# Destabilization of the Tumor-Inducing Plasmid from an Octopine-Type *Agrobacterium tumefaciens* Lineage Drives a Large Deletion in the Co-Resident At Megaplasmid

**DOI:** 10.1101/600809

**Authors:** Ian S. Barton, Thomas G. Platt, Douglas B. Rusch, Clay Fuqua

## Abstract

Bacteria with multi-replicon genome organizations, including members of the family *Rhizobiaceae*, often carry a variety of niche-associated functions on large plasmids. While evidence exists for cross-replicon interactions and co-evolution between replicons in many of these systems, remarkable strain-to-strain variation is also observed for extrachromosomal elements, suggesting increased genetic plasticity. Here, we show that curing of the tumor-inducing virulence plasmid (pTi) of an octopine-type *Agrobacterium tumefaciens* lineage leads to a large deletion in the co-resident At megaplasmid (pAt). The deletion event is mediated by a repetitive IS-element, IS66, and results in a variety of environment-dependent fitness consequences, including loss of independent conjugal transfer of the plasmid. Interestingly, a related and otherwise wild-type *A. tumefaciens* strain is missing exactly the same large pAt segment as the pAt deletion derivatives, suggesting a similar event over its natural history. Overall, the findings presented here uncover a novel genetic interaction between the two large plasmids of *A. tumefaciens* and provide evidence for cross-replicon integration and co-evolution of these plasmids.

## INTRODUCTION

Bacteria exhibit a diversity of genomic architectures and contain replicons that range from small, transient plasmids and megaplasmids, to chromids, to secondary and primary chromosomes (diCenzo and Finan 2017). Primary chromosomes are the largest replicons encoding core functions, whereas any replicating elements in addition to the primary chromosome are defined as secondary replicons. Secondary chromosomes are smaller than primary chromomes, but also carry core genes, and are thought to be derived directly from the primary chromosome. In contrast, chromids are defined as large replicons, (>350 kb) that also have essential genes, but appear to have arisen by horizontal gene transfer. Plasmids are replicons that carry accessory functions, but no truly essential genes, and if plasmids are larger than 350 kb are considered megaplasmids. Approximately 10% of bacteria possess multi-partite genomes and it is hypothesized that this arrangement may allow increased genome size, faster growth rate, coordinated gene regulation, and adaptation to novel niches (Chain *et al*. 2006; Slater *et al*. 2009; Galardini *et al*. 2013; diCenzo *et al*. 2014; diCenzo and Finan 2017). A multi-partite genome organization is likely maintained through selective pressures, but factors affecting replicon stability and their dynamic interactions are not well understood and often vary significantly between and within species.

Secondary replicons such as megaplasmids and plasmids confer a variety of niche-associated functions, such as antibiotic resistance, virulence, and symbiosis (Funnell and Phillips 2004; diCenzo and Finan 2017). While many of these replicons are dispensable in the laboratory, they often have profound positive and negative effects on bacterial fitness in a variety of environments and can affect growth rate and competition (Finan *et al*. 1986; Morton *et al*. 2013; diCenzo *et al*. 2014; Morton *et al*. 2014; diCenzo and Finan 2017), as well as numerous other phenotypes (Dougherty *et al*. 2014). Fitness consequences resulting from gain or loss of accessory genes and replicons are not typically measured, but likely depend on a combination of factors, including energetic demands of DNA replication, transcription and expression of specific costly functions, such as ABC transporters, as well as interactions between regulatory networks and pathways across replicons (Morton *et al*. 2013; Mauchline *et al*. 2014; Romanchuk *et al*. 2014; diCenzo and Finan 2017).

Interactions and genetic regulation between replicons are both a consequence of and a limitation to the coexistence of multiple replicons and have been observed in diverse bacterial species, including species of *Vibrio* (Heidelberg *et al*. 2000; Ramachandran *et al*. 2017) and *Sinorhizobium* (Ronson *et al*. 1987; Barnett *et al*. 2004; Bobik *et al*. 2006; Galardini *et al*. 2015; Pini *et al*. 2015; diCenzo *et al*. 2018). In most cases, chromosomal factors influence the regulation of secondary replicon factors, with limited regulation in the opposite direction (Ronson *et al*. 1987; Barnett *et al*. 2004; Bobik *et al*. 2006; Agnoli *et al*. 2012; Galardini *et al*. 2015; Pini *et al*. 2015; diCenzo *et al*. 2018), likely due to intolerance of integral pathway perturbation and the suppression of costly accessory functions required to stabilize secondary replicons. However, metabolic redundancy has been observed between rhizobial replicons (González *et al*. 2005; diCenzo and Finan 2017), suggesting that favorable interaction and pathway integration between primary and secondary replicons can occur. However, because the majority of organisms possessing multi-partite genomes exist and transition between multiple environmental reservoirs, and are often host-associated, the extent to which cross-replicon interactions occur and their consequences are often not determined.

Here, we characterize a cross-replicon genetic interaction in an octopine-type lineage of the plant pathogen *Agrobacterium tumefaciens,* causative agent of crown gall disease on plants (Van Larebeke *et al*. 1974; Watson *et al*. 1975; Escobar and Dandekar 2003). We observe that the process of curing the tumor-inducing virulence plasmid (pTi) is coincident with a large deletion in the co-resident At megaplasmid (pAt) of the octopine-type *A. tumefaciens* strain 15955. The deletion event is mediated by a repetitive IS-element and results in a variety of environment-dependent fitness consequences. Interestingly, there is evidence that the deletion identified here has also occurred in a related *A. tumefaciens* strain, suggesting that the deletion mechanism described may be common to a subset of strains in the *A. tumefaciens* genomospecies complex. Overall, the findings presented here indicate a novel and complex genetic interdigitization of these two secondary replicons in *A. tumefaciens* and provides mechanistic precedent for cross-replicon interactions between rhizobial plasmids.

## MATERIALS AND METHODS

### Reagents, media, strains, and growth conditions

All strains and plasmids used in this study are described in Table S1. Specific oligonucleotides are listed in Table S2, and all oligonucleotides used in the study are available upon request. Chemicals, antibiotics, and culture media were obtained from Fisher Scientific (Pittsburgh, PA), Sigma-Aldrich (St. Louis, MO), and GoldBio (St. Louis, MO) unless otherwise noted. Plasmid design and verification as well as strain creation was performed as described previously (Morton and Fuqua 2012a) with specific details included in the text. Oligonucleotide primers were ordered from Integrated DNA Technologies, Coralville, IA. Single primer extension DNA sequencing was performed by ACGT, Inc., Wheeling, IL. Plasmids were introduced into *E. coli* via transformation with standard chemically competent cell preparations and into *A. tumefaciens* via electroporation or conjugation (Morton and Fuqua 2012a).

**Table 1.** pAt15955 segment deletion and pTi15955 curing frequencies.

| Strain<br>(N) | Initial genotype |  | Ti curing conditions |  |
| --- | --- | --- | --- | --- |
|  | pAT | pTi | At deletion freq.<sup>a</sup> | Ti curing freq.<sup>b</sup> |
| WT IB118/IB126<sup>c</sup><br>(N, 684) | + | + | 0.71 ± 0.10 | 0.28 ± 0.08 |
| IB118 ΔIS66<br>(N, ~2500)<sup>d</sup> | + | + | <0.005 | ND<sup>e</sup> |
| IB129<br>(N, 261) | Δ270 | + (RI)<sup>f</sup> | NA<sup>e</sup> | 0.45 ± 0.17 |
| IB130<br>(N, 159) | + | + (RI)<sup>f</sup> | ND<sup>e</sup> | 0.13 ± 0.08 |
| IB118 ΔL<br>(N, 542) | Δ130 L | + | ND<sup>e</sup> | 0.27 ± 0.04 |
| IB118 ΔR<br>(N, 263) | Δ130 R | + | ND<sup>e</sup> | 0.42 ± 0.17 |
| IB118 ΔL+R<br>(N, 548) | Δ270 | + | NA<sup>e</sup> | 0.39 ± 0.07 |

<sup>a</sup>At deletion frequency determined as the ratio of *A. tumefaciens* curing plasmid transconjugants that have incurred the At plasmid deletion to total transconjugants.
<sup>b</sup>Ti curing frequency determined as the ratio of *A. tumefaciens* curing plasmid transconjugants that have lost the Ti plasmid to the total transconjugants. ~100% of isolates cured for pTi15955 have also incurred the pAt15955Δ270 deletion and only rare isolates have been recovered that retain the segment upon pTi15955 curing (text and Table S1).
<sup>c</sup>WT represents combined data from IB118 and IB126 for At deletion and Ti curing frequencies.
<sup>d</sup>From approximately 2500 colonies screened for *lacZ* on media with X-Gal with 500 secondarily screened by antibiotic resistance and Eckhardt gels, no transconjugants were found to have deleted the pAt segment.
<sup>e</sup>ND, not determined; NA, not applicable
<sup>f</sup>RI – indicates that pTi15955 was re-introduced by conjugation

*E. coli* was cultured in standard LB broth with or without 1.5% (w v^-1^) agar. Unless noted otherwise, *A. tumefaciens* strains were grown on AT minimal medium containing 0.5% (w v^-1^) glucose and 15 mM ammonium sulfate without added FeS0_4_ (ATGN) (Morton and Fuqua 2012b), or AT with 5% (w v^-1^) sucrose as the sole carbon source (ATSN). When required, appropriate antibiotics were added to the medium as follows: for *E. coli*, 100 μg ml^-1^ ampicillin (Ap), 50 μg ml^-1^ gentamicin (Gm), and 50 μg ml^-1^ kanamycin (Km); and for *A. tumefaciens*, 300 μg ml^-1^ Gm, 300 μg ml^-1^ Km, and 300 μg ml^-1^ spectinomycin (Sp). For competition experiments, *A. tumefaciens* 15955 strains were grown in replete carbon (standard ATGN), limiting carbon (ATGN with 0.018% (w v^-1^) glucose) or virulence-inducing media (Modified ATGN, replacing AT buffer with 50 μM phosphate buffer (NaH_2_PO_4_, Na_2_HPO_4_, pH 5.7) with 0.02M 2-N-[morpholino]ethanesulfonate (MES), and supplementing with 200 μM acetosyringone (Morton and Fuqua 2012b). Octopine utilization was evaluated by growing *A. tumefaciens* 15955 in AT media modified to contain octopine (3.2 mM) as the sole carbon and nitrogen source (ATO).

Precise deletion of IS66 was achieved using a previously described method (Morton and Fuqua 2012a). Fragments ranging from 500-750 bp and corresponding to sequences upstream (P1 and P2) and downstream (P3 and P4) of the gene to be deleted were amplified using Phusion polymerase (New England Biolabs, Ipswich, MA). Primers P2 and P3 were designed with 5’ sequences with reverse complementarity to each others’ 3’ proximal sequence. The complementary sequence on these two primers facilitated splicing by overlapping extension (SOE) of the two PCR products, as previously described (Morton and Fuqua 2012a). Both flanking sequences were amplified and gel purified. The two purified products were then used as both templates and primers in a second PCR reaction, generating the final spliced product, which was ligated into pGEM-T easy (Promega) and sequenced to ensure proper splicing. The deletion construct was then excised using restriction enzymes and ligated into the suicide vector pNPTS138 cleaved with compatible restriction enzymes. The pNPTS138 plasmid confers Km resistance (Km^R^) and sucrose sensitivity (Suc^S^). Derivatives of pNPTS138 were introduced into *A. tumefaciens* 15955 by mating with *E. coli* S17-1/λpir carrying the appropriate construct. The ColE1 origin of pNPTS138 does not replicate in *A. tumefaciens*, necessitating single-crossover integration into the chromosome to obtain Km^R^ transformants. Plasmid integration was confirmed by patching onto ATGN-Km and ATSN to identify Suc^S^ derivatives. Excision of the integrated plasmid was then mediated by growing overnight cultures of Suc^S^ Km^R^ derivatives and plating dilutions onto ATSN. Plasmid excision was confirmed by patching Suc^R^ clones onto ATSN and ATGN-Km to identify Km^S^ derivatives. Correct deletion of the target gene was confirmed via diagnostic PCR using primers (P5/P6) flanking the deletion site.

Marked derivatives of *A. tumefaciens* 15955 were created using an approach similar to the creation of in-frame deletions (above) with minor alterations. P2 and P3 were designed to have complementarity in the form of a multiple cloning site, consisting of five restriction sites. After splicing of the PCR-amplified product into pNPTS138, the restriction sites were used to fuse in different screenable and selectable markers. Plasmid excision in Km^S^ derivatives of Suc^R^ clones were confirmed similarly, and retention of the screenable or selectable markers was verified by sequencing of PCR products.

### Whole genome sequencing and bioinformatic analysis

Whole genome sequencing and assembly was performed at the Center for Genomics and Bioinformatics, Indiana University, Bloomington, IN. WT *A. tumefaciens* 15955 was sequenced using a combination of 454 and Illumina NextSeq platforms and assembled *de novo*. Sequencing of *A. tumefaciens* 15955 derivatives was performed using Illumina NextSeq 300 with 150 bp paired-end read libraries. Raw reads were trimmed using Trimmomatic ver. 0.33 (Bolger *et al*. 2014) and bowtie2 ver. 2.2.6 (Langmead and Salzberg 2012) was used to map trimmed reads to the assembled WT *A. tumefaciens* 15955 genome sequence (above). Mapping efficiency was >99% and concordantly mapped reads had an average insert size of ∼500 bp with a normal distribution. Average coverage across the genome was at least 20X. Variation between WT *A. tumefaciens* 15955 and derivatives were determined using breseq ver. 0.24rc6 (Deatherage and Barrick 2014). Genome alignments and comparisons were performed with mummer and mummerplot packages of MUMmer 3.23 (Kurtz *et al*. 2004) and Artemis Comparison Tool (ACT) (Carver *et al*. 2005) visualizations of TBLASTX (https://blast.ncbi.nlm.nih.gov/Blast.cgi) data. Over the course of this project a group from the Academia Sinica in Taiwan deposited a complete genome sequence for the wild type *A. tumefaciens* 15955 into GenBank (Accession numbers; Bioproject PRJNA494475; Circular chrom. NZ_CP032917.1; Linear chrom. NZ_CP032918.1; pAt15955 NZ_CP032919.1; pTi15955 NZ_CP032920.1). This sequence is in near-perfect agreement with our *de novo* wild type sequence, and we refer to the coordinates of this deposited sequence where relevant.

### Southern blotting

A ∼500 bp DIG-labeled DNA probe (Sigma-Aldrich) specific to IS66 was generated with PCR (1:6 ratio dig-UTP:dTTP) using primers IBP102/IBP103 and used to probe 2.5 μg restriction digested gDNA (*Sph*I, *Eco*RI, *Bam*HI) that had been electrophoresed on a 0.75% agarose gel (2h, 50V) and transferred to a charged nylon membrane via capillary blotting under alkaline conditions, as described previously (Fuqua and Weiner 1993). Restriction digests were performed at 37°C for 2 hours. Hybridizations and detection of the DIG-labeled DNA probe were performed according to manufacturer’s suggestions (Sigma-Aldrich). Blots were visualized using a Bio-Rad Chemidoc system.

### pTi15955 curing and detection of pAt15955 segment loss

Curing of pTi15955 was performed as described by our lab previously (Morton and Fuqua 2012a) with slight modifications. The curing plasmid is similar to the pBBR1-type curing plasmids recently described (Yamamoto *et al*. 2018), containing two origins capable of replication in *Agrobacterium*, a Km^R^ cassette, and the counterselectable *sacB* marker for removal of the curing plasmid. Briefly, pSRKKm was digested with *Bam*HI (NEB) according with NEB guidelines, followed by purification using the E.Z.N.A. Cycle Pure Kit from Omega Bio-tek. The digested and purified pSRKKm backbone was then used in a three-part NEBuilder isothermal assembly reaction (New England Biolabs) with *repABC* (pTi15955) and *sacB* PCR products that were amplified from *A. tumefaciens* 15955 gDNA or purified pNPTS138, respectively, using primer pairs IBP200/IBP201 and IBP202/203 to create pIB301. NEBuilder reactions were performed as specified by the supplier using a 1:2 vector to insert ratio. pIB301 was confirmed using a combination of PCR and sequencing.

The curing plasmid pIB301 was introduced into *A. tumefaciens* 15955 backgrounds via conjugation from *E. coli* S17-1/λpir donor and *A. tumefaciens* 15955 transconjugants were identified through selective plating. The pIB301 plasmid carries broad host range replication origin (pBBR-type) and also containing a copy of the pTi15955 replication origin (*repABC*), a selectable kanamycin resistance marker (Km^R^), and a counterselectable marker (*sacB*). The plasmid is conjugated into *A. tumefaciens* 15955 strains from an *E. coli* donor and the resulting Km^R^ transconjugants were screened for the loss of the Ti plasmid. Ti plasmids generally have a low copy number under standard laboratory conditions (Suzuki *et al*. 2001; Pappas 2008; Platt *et al*. 2014).

To increase the efficiency by which both curing of pTi15955 and loss of the pAt15955 segment could be diagnosed, screenable and selectable marker cassettes were introduced into the *A. tumefaciens* 15955 genome within the pAt15955 segment (pos. 364,474) and on pTi15955 (pos. 17,447) using recombinational mutagenesis strategies described previously (Morton and Fuqua 2012a). A *lacZ*/Sp^R^ cassette and a *gusA*/Gm^R^ cassette were introduced between convergently oriented genes within the pAt15955 segment and on pTi15955, respectively, to create *A. tumefaciens* IB118. Upon introduction of the curing plasmid, frequencies for loss of the pAt15955 segment and pTi15955 curing were determined through a combination of blue/white screening on X-Gal (β-galactosidase, LacZ indicator) or X-Gluc (β-glucuronidase, GusA indicator), followed by confirmation with Sp^S^ or Gm^S^ screening, respectively, and validated by PCR and Eckhardt gel analysis. Loss of LacZ or GusA activity and Sp or Gm sensitivity in these reporter strains reported on pAt15955 segment loss or pTi15955 curing, respectively, with 100% fidelity (see Table 1 for representative use of these marked derivatives).

Screening for pTi15955-cured isolates and loss of the pAt15955 segment was performed using loss of *gusA* or *lacZ* activity on 5-bromo-4-chloro-3-indolyl-β-glucuronic acid (X-Gluc) or 5-bromo-4-chloro-3-indolyl-β-D-galactopyranoside (X-Gal) as well as Gm or Sp sensitivity, respectively, and/or a combination of Eckhardt gel electrophoresis (Eckhardt 1978) and PCR. Eckhardt gel electrophoresis was performed according to a modified protocol (Personal communication, J. Griffitts, Brigham Young University, Provo, UT). Additionally, the presence of pTi15955 was also evaluated using octopine auxotrophy screening.

Curing frequencies were statistically analyzed by pairwise, two tailed (heteroscedastic) Student’s *t*-tests in MS-Excel.

### Ti plasmid reintroduction and conjugation assays

pTi15955::*gusA/*Gm^R^ was reintroduced into Sp^R^ *A. tumefaciens* 15955 backgrounds (IB128/IB129) through conjugation with IB123 (At+Ti+::*gusA*/Gm^R^) as a donor in ATGN media supplemented with octopine [3.25 mM]. Briefly, donor and recipients were grown in ATGN supplemented with appropriate antibiotics to OD_600_ 0.6 and 100 µL of donor and recipient were combined and spotted onto sterile 0.2 µm cellulose acetate filter discs placed on ATGN media supplemented with octopine (ATGNO [3.25 mM]). After incubation (18 h, 28°C), the spots were resuspended in ATGN and plated onto ATGN supplemented with antibiotics to select for transconjugants. Transconjugants were confirmed using Eckhardt gel electrophoresis and PCR.

Conjugation assays were performed as described previously (Heckel *et al*. 2014) with alterations. Briefly, donors and recipient were grown and normalized to OD_600_ 0.6, mixed 1:1, and 100 µL was spotted onto 0.2 µm cellulose acetate filter discs placed on ATGN supplemented with and without octopine [3.25 mM]. Following incubation (18h, 28°C), matings were suspended in liquid ATGN, serially diluted, and spotted onto media supplemented with appropriate antibiotics. Conjugation efficiency was determined as the ratio of transconjugants per output donor.

### Truncation of pAt15955 segment using Cre-lox

The *loxP* site (5’ GCGGTCGAC<u>ATAACTTCGTATA</u>*ATGTATGC*<u>TATACGAAGTTAT</u>CATATGGCG 3’, 18 bp inverted sequences are underlined, and the 8bp spacer is italicized) was introduced into four locations within pAt15955::*lacZ*/Sp^R^ of IB118 in a pair-wise fashion to truncate either half of, or the entire pAt15955 segment upon transient expression of the Cre recombinase. To allow Cre-mediated deletion the left half of the At plasmid segment (denoted L), *loxP* sites were introduced in a tandem, direct orientation into positions L and CL that are interior to the flanking IS66 element and *lacZ*Sp^R^ marker in IB118, respectively, to create IB118 *lox*L (Figure S5 and Table S1). Deletion of right half (denoted R) of the segment was done similarly, except *loxP* sites were integrated at positions CR and R to create IB118 *lox*R. *loxP* sites were integrated at positions L and R to create IB118 *lox*L+R and allow deletion of the entire segment. Confirmation of genomic integration of each *loxP* site was done using PCR with primers specific for each integration site. Transient expression of the Cre recombinase was accomplished through introduction of the pCRE1-Ap suicide vector (Bailey and Manoil 2002) into the appropriate *A. tumefaciens* 15955 recipient (IB118 *lox*L, IB118 *lox*R, or IB118 *lox*L+R) via mating with *E.coli* S17-1/λpir pCRE1-Ap as described (Bailey and Manoil 2002) with minor alterations. After mating of donor and recipients on LB agar, resuspensions were plated onto LB with or without Sp supplementation for deletion of the left and right half or entire segment, respectively. Confirmation of Cre-mediated deletions were confirmed using Eckhardt gel electrophoresis and PCR with primers specific for each new junction site created through Cre-mediated deletion.

### Competition experiments

Competition experiments were performed as described previously with minor alterations (Morton *et al*. 2013). Competitor strains were grown in ATGN with or without antibiotic to an OD600nm of 0.6-0.8, washed 3 times in ddH_2_0, and mixed 1:1 at an OD600nm of 0.01 in 2 mL of media. When competitions reached an OD600 nm of 0.6-0.8, 0.4-0.6, or 0.2-0.4 for replete carbon, limiting carbon, and virulence-inducing media, respectively, the competitions were serially diluted and spotted onto ATGN with and without antibiotic to enumerate each competitor, as well as sub-cultured (1:100) into the same media. Competitions were carried out for a total of nine passages and each was performed 3 times with reciprocally-marked strains for a total of 6 replicates. Relative fitness was estimated as the number of doublings by each competitor over each passage during the course of the experiment, as described previously (Lenski 1988; Morton *et al*. 2013). Passaging intervals, such as P0-P9, thusly denote cumulative doublings over every passage.

### Data and reagent availability

The *A. tumefaciens* 15955 genome sequence is available through GenBank Accession No. PRJNA494475, and all coordinates for this genome sequence cited in the manuscript correspond to this sequence. All reagents, including strains, plasmids, and oligonucleotides are available upon request Supplementary information, including figures, tables and references for this study have been deposited as a single file in GSA figshare (filename: Supplementary_Information_G3_Revised).

## RESULTS

### Large deletion in pAt15955 upon pTi15955 curing

Several on-going studies rely on curing of the Ti plasmid from *A. tumefaciens*, using a plasmid incompatibility, eviction process (Uraji *et al*. 2002; Platt *et al*. 2012a; Yamamoto *et al*. 2018). As a matter of practice, the plasmid profiles of putatively cured derivatives are analyzed by Eckhardt gel electrophoresis to visualize their large plasmid content (see Materials and Methods). Eckhardt gel analysis of *A. tumefaciens* 15955 revealed two discrete bands, reflecting the presence of both an At plasmid (pAt15955, large band) and Ti plasmid (pTi15955, smaller band) (Figure 1). Several derivatives of *A. tumefaciens* 15955 were cured of their Ti plasmid using the incompatibility-eviction approach. Strikingly, it was observed that plasmid profiles of these cured derivatives of *A. tumefaciens* 15955 incurred multiple changes, where bands corresponding to the Ti and At plasmids were replaced by a band that migrated faster than pAt but more slowly than pTi (Figure 1). This change in plasmid profile was observed for virtually all of the Ti plasmid-cured derivatives from multiple experiments. Interestingly, Eckhardt gel analysis of *A. tumefaciens* SA122, a 15955 derivative cured by a completely different approach using chemical and temperature stress in a different laboratory years earlier revealed the same pattern as the derivatives we had generated (Figure 1, and see Materials and Methods) (Uraji 2002; Yamamoto *et al*. 2018).

**Figure 1:**
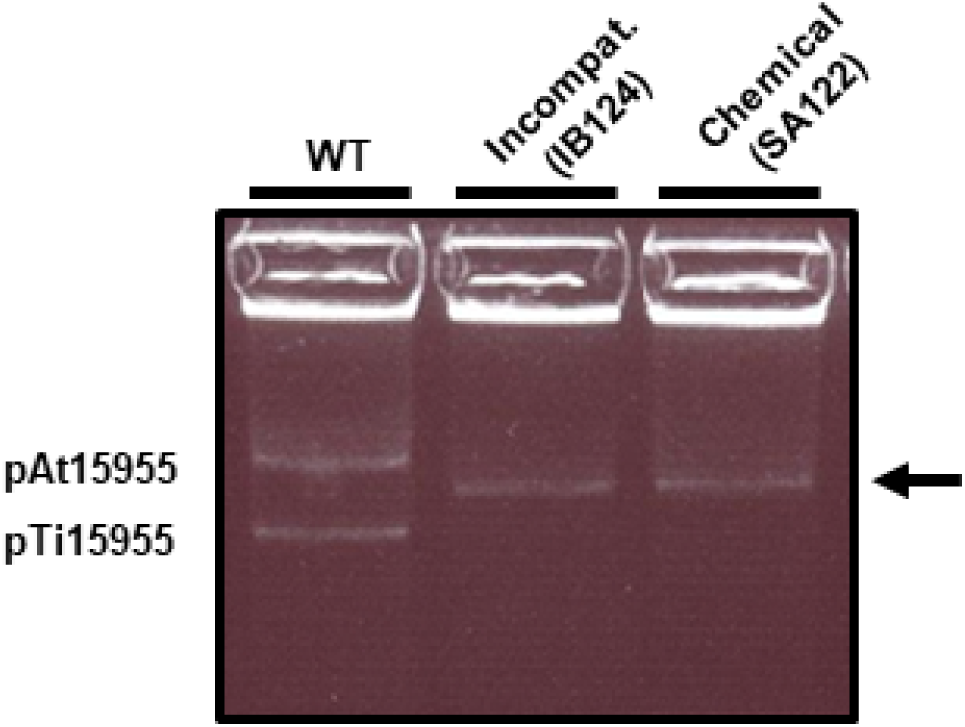
Multiple changes in plasmid profile of *A. tumefaciens* 15955 upon curing of pTi15955. Eckhardt gel electrophoresis analysis of WT *A. tumefaciens* 15955 and two derivatives of *A. tumefaciens* 15955, cured of the Ti plasmid (pTi15955) using either chemical (SA122) or plasmid incompatibility methods (IB124). Positions of pAt15955 and pTi15955 in WT *A. tumefaciens* 15955 are shown. Arrow indicates single band present in pTi15955-cured derivatives.

To confirm loss of pTi15955, as well as diagnose other genomic changes in these strains underlying the changes observed using Eckhardt gel electrophoresis, whole genome sequencing was conducted on WT *A. tumefaciens* 15955 and pTi15955-cured derivatives. WT *A. tumefaciens* 15955 assembled into 13 scaffolds, the first four corresponding to the circular chromosome (scaffold00001, 2.836 Mbp), linear chromosome (scaffold 00002, 2.082 Mbp), pAt (scaffold00003, 0.816 Mbp), and pTi (scaffold00004, 0.192 Mbp). Nine other smaller scaffolds did not assemble with the four primary scaffolds and ranged in size from 16.045 Kbp (scaffold00005) to 2.419 Kbp (scaffold00013). The sequencing confirmed loss of pTi15955 in cured derivatives but also identified a large (270,949 bp) deletion in the pAt15955 (Figure 2) that was consistent between multiple pTi15955-cured isolates (Figure 2 and data not shown). From this, we concluded the single band present in pTi15955-cured isolates upon Eckhardt gel electrophoresis is the truncated pAt15955 (denoted as pAt15955Δ270) (Figure 1). The *A. tumefaciens* 15955 genome sequence recently deposited in GenBank database (Bioproject PRJNA494475) does not have this deletion, and matches our *de novo* generated wild type sequence very closely.

**Figure 2:**
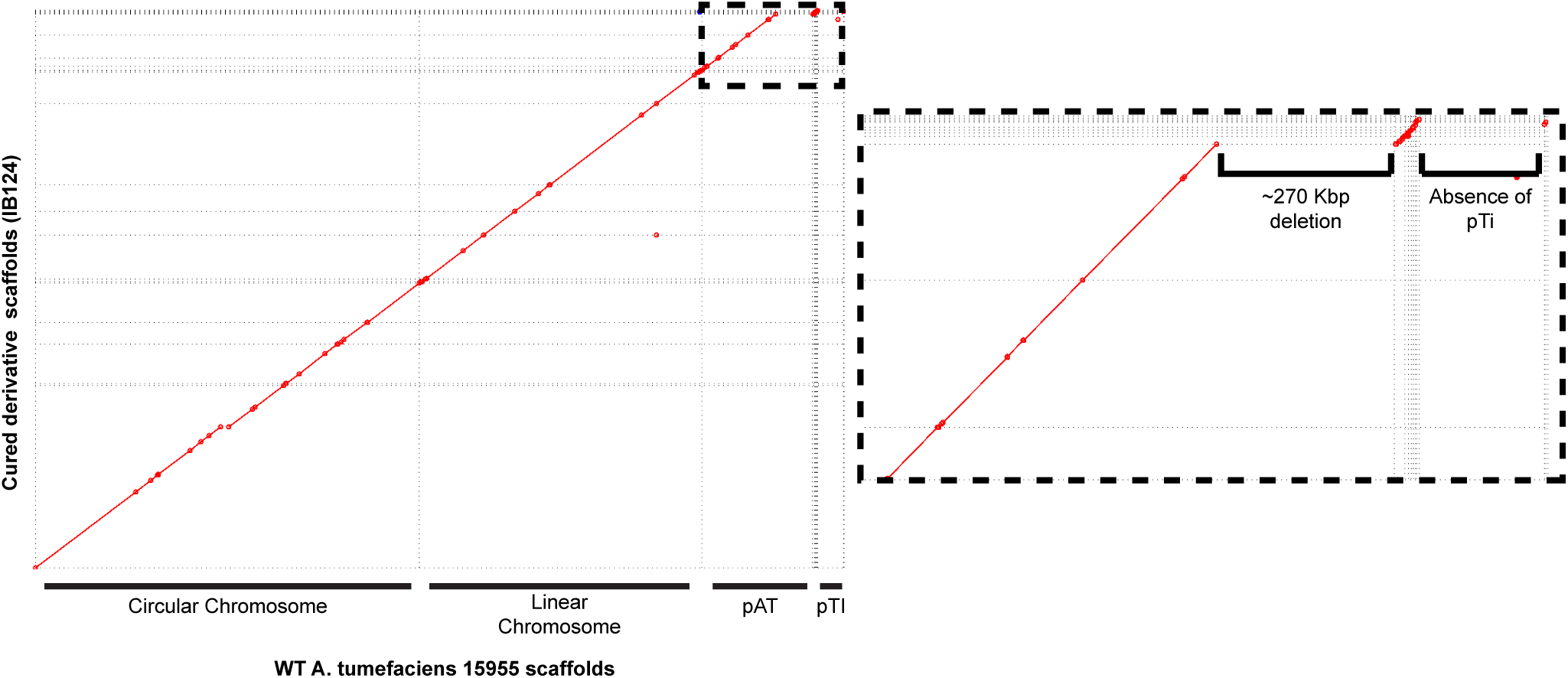
Whole genome sequencing reveals co-incident pAt15955 deletion upon pTi15955 curing. Mummerplot visualization of MUMmer 3 alignment of genome assembly scaffolds from WT 15955 (X axis) and a pTi15955-cured derivative (IB124, Y axis). Matching sequence between assemblies is indicated with diagonal red line segments. Sequence corresponding to each replicon is indicated. Inset (dashed) highlights large (270 Kbp) deletion of pAt15955 and absence of pTi15955 in IB124.

### Evaluation of curing and deletion frequencies suggests these processes are linked but independent

We have observed two separate, large scale genomic events, Ti plasmid curing and the large deletion incurred in pAt15955, occurring coincidentally. To characterize the connection between these events, the correlation between pTi15955 curing and pAt15955 segment loss was determined under pTi15955 curing conditions. High efficiency incompatibility-based curing strategies were used to determine the frequency of both events (Uraji 2002; Yamamoto *et al*. 2018) as detailed in the Materials and Methods. Plasmids with the pTi15955 replication origin were introduced by conjugation into *A. tumefaciens* 15955 derivatives. Transconjugant populations were subsequently screened for loss of the Ti plasmid and the pAt deletion event. We genetically marked the Ti plasmid and the 270 kb segment of the At plasmid with a *lacZ/*Sp^R^ and a *gusA*/Gm^R^, respectively, to generate *A. tumefaciens* 15955-IB118 to facilitate initial screening for curing and deletion events (described in Materials and Methods).

Using *A. tumefaciens* 15955-IB118 derivative with the *lacZ*/Sp^R^ and a *gusA*/Gm^R^ markers as a recipient for the curing plasmid, it was determined that ∼20% of Km^R^ transconjugants have lost pTi15955 (Table 1). Consistent with previous analyses of pTi15955-cured isolates (Figure 1), nearly 100% of pTi15955-cured isolates also incur loss of the pAt15955Δ270 segment (Table 1). However, ∼71% of curing plasmid transconjugants incur loss of the segment, indicating that it is deleted at a higher frequency than loss of the pTi15955 plasmid under these conditions (Table 1). Taken together, the correlation suggests that curing conditions stimulate pTi15955 curing and pAt15955 segment loss, that the events are separable, and that loss of the pAt15955 may precede, and/or possibly even limit, pTi15955 curing.

We also explored whether more general stress conditions could induce the observed pAt15955 deletion. Multiple individual isolates of *A. tumefaciens* 15955 isolates grown under standard conditions, antibiotic exposure, heat shock (37°C), and virulence-inducing conditions (acidic pH and plant phenolics) were examined for the deletion. None of the isolates tested (∼3.0 x 10^8^) had incurred the deletion suggesting that it is specifically induced by the curing process, although independent of the exact curing mechanism.

### *A. tumefaciens* Ach5, a related octopine-type strain that lacks the segment deleted from pAt15955

Strains belonging to the *A. tumefaciens* species complex have been classified based on their genome sequences into several genomospecies (Mougel *et al*. 2002), and *A. tumefaciens* 15955 falls into the G1 genomospecies (Henkel *et al*. 2014; Huang *et al*. 2015). *A. tumefaciens* Ach5 is another G1 genomospecies closely related to *A. tumefaciens* 15955 and carrying a an octopine-type Ti plasmid. Analysis of the Ach5 genome sequence (GenBank Access. No. Bioproject PRJNA278497) reveals that the pAtAch5 megaplasmid plasmid it harbors is missing the exact same large segment as we have observed to delete in pAt15955 under Ti plasmid curing conditions, with nearly identical sequence flanking this region (Figure S1) (Henkel *et al*. 2014; Huang *et al*. 2015).

### The pAt15955 deletion event is mediated by flanking IS66 elements

Analysis of whole genome sequencing data identified three copies of an IS element, IS66, within the *A. tumefaciens* 15955 genome, one on the linear chromosome (pos. 28,096-30652) and two on pAt15955 (IS66_At_L, pos. 508,310-510,865 and IS66_At_R, pos. 237,361-239,916), flanking the 270 Kbp segment that is deleted upon elicitation of Ti plasmid curing conditions (Figure S2). IS66 represents a family of IS elements conserved within a wide variety of bacteria, but was first identified on an octopine-type Ti plasmid within *A. tumefaciens* (Machida *et al*. 1984; Han *et al*. 2001; Gourbeyre *et al*. 2010). The element consists of three open reading frames, *tnpABC*, the latter of which is thought to encode an active transposase and is the most conserved across the IS66-family. The three ORFs are bordered by distinct direct and indirect repeats (Figure S2).

Repetitive sequences, such as IS-elements, are notoriously difficult to correctly assemble in next generation sequencing data. We aimed to confirm and enumerate the IS66 elements within *A. tumefaciens* 15955 and its deletion derivative using a Southern blot approach. A 515 bp DNA fragment specific for *tnpC* was generated by PCR (primers IBP102 and IBP103) and used to probe restriction enzyme-digested genomic DNA (*Sph*I, *Eco*RI, and *Bam*HI) from *A. tumefaciens* 15955 and its derivatives. In WT *A. tumefaciens* 15955 gDNA, three distinct bands were present in restriction digests, confirming that *A. tumefaciens* 15955 contains three copies of IS66 (Figure 3A). Fragment sizes of detected bands are consistent with two IS66 elements flanking the pAt15955 segment (Figure 3D). Strikingly, only two visible bands are detectable for gDNA from the IBE13A derivative, deleted for the pAt15955Δ270 plasmid segment, indicating loss of one IS66 element during the deletion event (Figure 3B). One of the two fragments matches a fragment in the wild type, and the second for the *Sph*I and *Eco*RI digests does not match either of the other two restriction fragments in the wild type. The *Bam*HI digest of the deletion mutant generates a predicted new fragment very close in size to the previous two. These observations are consistent with loss of one IS66 element and retention of a single copy of IS66 on pAt15955 at the site of deletion, and correlates with predictions from our own sequence analysis (Figure 3D) and examination of the GenBank 15955 accession.

**Figure 3:**
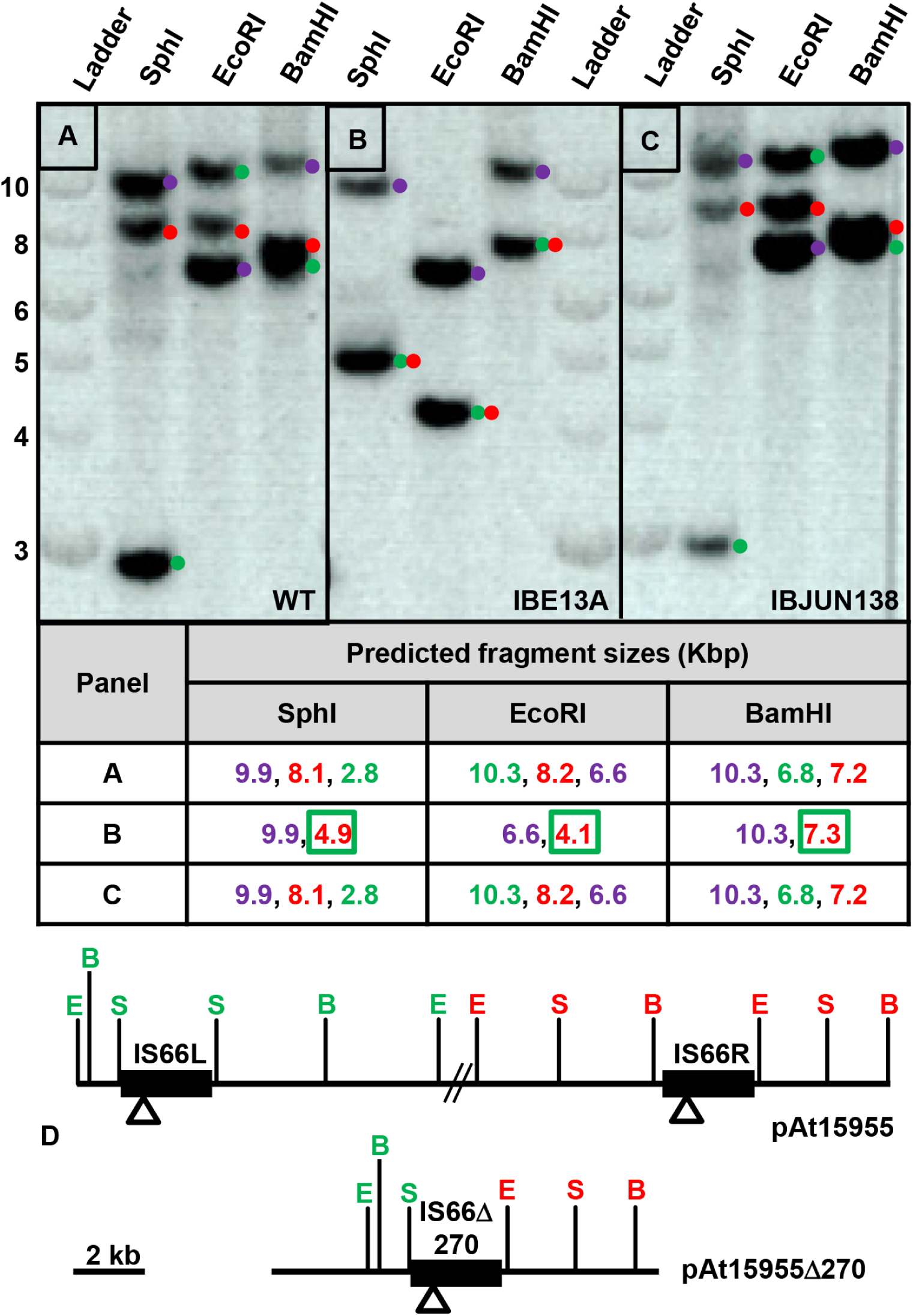
Loss of one IS66 element upon deletion of ∼270 Kbp pAt15955 segment. Southern blot analysis of gDNA from WT *A. tumefaciens* 15955 (**A**) and two derivatives that have either incurred loss of the pAt15955 segment (IBE13A, **B**) or pTi15955 (IBJUN138, **C**) using three restriction enzyme digests that do not cleave within IS66 (SphI, EcoRI, and BamHI). Ladder shown is NEB 1 kb DNA ladder. Numbers on Y-axis indicated ladder sizes in Kbp. Prediction of IS66-containing restriction fragment sizes for digests in A, B, and C – color coding matches the dots adjacent to each band; Blue, IS66 on the linear chromosome; Green, IS66L; Red IS66R; New restriction fragments generated by the deletion are red text boxed green (**D**) Restriction site locations in the intact pAt15955 plasmid and the pAt15955Δ270 derivative. S, SphI; E, EcoRI, B, BamHI. Color coding of restriction sites matches predicted fragments and the annotated blot image. Open triangles represent the location of the probe fragment used for the hybridizations, and the double hash marks are the remaining 270 kb deleted from pAt15955..

While nearly all *A. tumefaciens* 15955 curing plasmid transconjugants that lose pTi15955 also lose the pAt15955 segment, we have recovered several rare isolates that have been cured of pTi15955 but do not carry the deletion, including derivative IBJUN138. Southern analysis of IBJUN138 did not deviate from the pattern observed for *A. tumefaciens* 15955 (Figure 3C), confirming our sequence assemblies suggesting that IS66 does not reside on the Ti plasmid in this strain and rather is on the linear chromosome, in contrast to previous reports for IS66 on octopine-type Ti plasmids in other *A. tumefaciens* strains (Machida *et al*. 1984).

Retention of a single IS66 element following the deletion event suggests homologous recombination as the primary mechanism for segment deletion. Recombination-mediated deletion of the pAt15955 segment would require that both elements be in the same orientation (Petes and Hill 1988). An inverse PCR approach was used to confirm that both IS66 elements flanking the pAt15955 segment in WT *A. tumefaciens* 15955 exist in a tandem orientation (Figure S4A). Based on predicted restriction fragment sizes surrounding each IS66 element in WT *A. tumefaciens* 15955, inverse PCR products support the direct orientation of both elements (Figure S4B and data not shown), also in agreement with the sequence assemblies.

As repetitive IS-elements are common substrates for homologous recombination, it seemed likely that the two IS66 elements on pAt15955 were responsible for the observed deletion during pTi15955 curing. To examine the role for both IS66 elements in mediating the pAt15955Δ270 deletion, the left copy of the IS66 element (IS66-At-L) was deleted from pAt15955 within the reporter strain IB118. The resulting strain (IB118ΔIS66) was subjected to pTi15955-curing conditions identical to those used for the other curing experiments and evaluated for pAt15955 segment loss. Derivatives lacking the pAt15955Δ270 deletion segment were undetectable in IB118ΔIS66 under pTi15955 curing conditions (∼2500 colonies screened, Table 1), consistent with the requirement for the two elements to drive homologous recombination. Interestingly, *A. tumefaciens* Ach5 also has a single copy of IS66 at the deletion site in pAtAch5, suggesting that IS66 may have mediated a similar large deletion in this strain (Figure S3). Insertion of many transposable elements creates a short duplication at the site of insertion, and this is reported to be 8 bp for IS66 (Machida et al. 1984). Unique 8 bp duplications flank all three of the IS66 elements in the 15955 sequence (IS66_Lin, AGAACACG; IS66_At_L, CCTTGATC; IS66_At_R, CGTTCCGG). In contrast, for the pAt15955Δ270 deletion derivative the 8 bp sequences immediately flanking the left and right ends in the single remaining At plasmid IS66 element match the outside flanking sequence from the IS66_At_L and IS66_At_R, respectively (Fig. S2). Similarly, the 8 bp flanking each end of IS66 in pAtAch5 are different from each other (CCGGAACG and CCTTGATC) and match those flanking the IS66 pAt15955Δ270. These observations are again consistent with homologous recombination between IS66 elements driving the deletion events in both the ancestor of pAtAch5 and in pAt15955.

### Genetic analysis of pTi15955 stabilization by the deleted segment in AtΔ270

Our observations suggest a step-wise mechanism in which loss of the pAt15955 segment usually precedes pTi15955-curing (Table 1). However, we were able to obtain a rare derivative in which the Ti plasmid was cured, but the pAt15955 plasmid did not have the Δ270 deletion (IBJUN138). It seemed likely that this derivative had incurred mutations which abolished the requirement for loss of the At plasmid segment before Ti plasmid curing. However, whole genome sequence analysis of IBJUN138, did not reveal mutations within the deleted segment and none that would readily explain retention of this pAt segment. We therefore used the IBJUN138 strain that had retained the intact At plasmid but was cured for the Ti plasmid, to examine the impact of the retained At plasmid genes on curing frequencies. The *gusA*/Gm^R^-marked pTi15955 plasmid was reintroduced by conjugation back into two pTi15955-cured derivatives with a chromosomal Sp^R^ marker (these derivatives do not carry the pAt15955 *lacZ*/Sp^R^ marker), one that had incurred loss of the pAt15955 segment (IB129) and another that had not (IB130, a derivative of IBJUN138) (Table S1 and Methods). These strains were then subjected to pTi15955-curing conditions and evaluated for Ti plasmid loss. The reintroduction derivative, IB130, appeared to have a modest decrease in pTi15955 curing relative to other strains, and the IB129 strain that had already incurred the pAt15955 deletion was also slightly elevated in pTi curing frequency (Table 1). However, none of these measured differences were statistically significant to a P-value <0.1 (by Students *t*-test). The observed trends suggested that the presence of the pAt15955Δ270 segment might weakly stabilize the Ti plasmid, but the differences measured were too variable to allow an unequivocal interpretation.

We also sought to determine whether a specific genetic locus within the deleted segment might impact curing efficiency. To test this hypothesis, the derivative carrying the marker in the deleted segment (IB118) was truncated for 138.8 kb on the left side (IB118 ΔL, pos. 504,337-365,531) or 106.7kb on the right side (IB118 ΔR, pos. 354,236-247,489) portion of the deleted segment, or the entire (IB118 ΔL+R) segment, using a Cre/*lox*-driven deletion approach (Figure S5, Table S1, and Methods). Strains truncated with Cre/*lox* retain both flanking IS66 copies and retain the *lacZ*Sp^R^ markers within the deletion interval, except for IB118ΔL+R, in which the *lacZ*Sp^R^ markers are deleted but flanking IS66 elements are still retained (Figure S5). These engineered deletion derivatives were subjected to pTi15955-curing conditions, and evaluated for pTi15955-curing frequencies. Truncation of part, or the entire pAt15955 segment again had modest effects on pTi15955 stability (Table 1). The two derivatives deleted for either the entire element or its right half exhibited a measureable increase in pTi15955 curing frequency, whereas the deletion of the left half of this segment was very similar to the wild type frequency (Table 1). However, these trends were not statistically significant to a level of P-values <0.1 (by Students *t*-test). This suggests that genetic elements within the right half of the pAt15955 segment might influence pTi15955 stability, but this effect is weak (Table 1). Similar Ti plasmid curing frequencies were observed for the isolated derivative deleted for the pAt15955 segment (IB129) with the single IS66 element and the engineered derivative lacking the genes on the segment, but retaining the tandem IS66 elements (IB118 ΔL+R), suggesting that the tandem IS66 elements alone do not influence pTi15955 stability (Table 1).

### The pAt15955Δ270 segment encodes conjugation machinery and conjugal transfer of pAt15955 is octopine-inducible

Analysis of whole genome sequence data from WT *A. tumefaciens* 15955 identified conjugal transfer genes within pAt15955 that are homologous to genes on pTi15955 (Figure S6 and data not shown). Interestingly, unlike pTi15955 and other Ti plasmids, pAt15955 *trb* and *traI* are not directly divergent from *repABC* and are rather encoded within the pAt15955Δ270 segment (Figure S6). Thus, strains deleted for the pAt15955 segment are lacking *trb* and *traI* genes, which are essential for conjugation in Ti plasmids with similar systems (Li *et al*. 1998; Li *et al*. 1999), and so they may be incapable of conjugation. To test this, WT *A. tumefaciens* 15955 and derivatives that lack the pAt15955 segment, pTi15955, or both, were marked with Km^R^ and used in conjugation assays with the ERM52 recipient, a plasmidless (At^-^Ti^-^) *A. tumefaciens* C58 derivative marked with Sp^R^ (Figure 4, Methods). In standard growth conditions (ATGN), low level At plasmid conjugation was observed for the marked wild type (IB125), and this is abolished in the deleted derivative (AtΔ270 Ti^+^) lacking the pAt15955 *trb* genes (IB131). (Figure 4). Conjugation of the Ti plasmid in octopine-type *A. tumefaciens* strains is strictly regulated by the presence of octopine and a complex quorum sensing process (Piper *et al*. 1993; Fuqua and Winans 1994; Hwang *et al*. 1994; Gordon and Christie 2014). Addition of octopine stimulates conjugal transfer frequency by two orders of magnitude in the At^+^Ti^+^ donor (IB125). Surprisingly octopine also stimulated significant pAt15955 conjugation in the Ti^+^AtΔ270 deletion strain (Figure 4). The Ti^-^ AtΔ270 derivative (IB132) did not detectably conjugate, and conjugation in the At^+^Ti^-^ strain (IB133) was the same as the uninduced At^+^Ti^+^, but was not stimulated by octopine addition. These findings suggest that the pAt15955 can drive its own low-level conjugation, dependent on conjugal transfer genes in the deleted segment (*trb* and *traI*). Octopine addition stimulates pAt15955 conjugation and this is dependent on the Ti plasmid, and can even overcome the absence of the pAt15955Δ270 segment.

**Figure 4:**
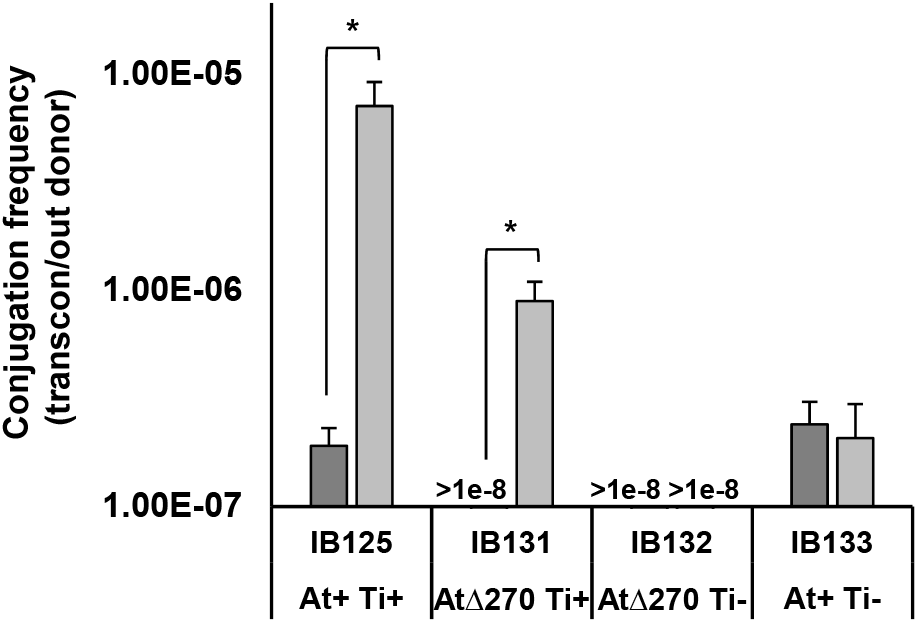
Octopine-inducible pAt15955 conjugation is pTi15955-dependent and independent of pAt15955-encoded *trb* machinery. pAt15955 conjugation frequencies of marked (Km^R^) derivatives of *A. tumefaciens* 15955 (donors) into a plasmidless recipient, ERM52, under standard growth (ATGN, dark gray) and octopine-containing (ATGNO, hatched, 3.25 mM) conditions. Donor strains shown are WT (IB125) or derivatives deleted for pAt15955 segment (IB131), pTi15955 (IB133), or both (IB132). Conjugation frequencies reported as transconjugants per output donor (see Materials and Methods). The * denotes a p-value <0.03.

One explanation for pTi15955-dependent, octopine-induced transfer of pAt15955Δ270 is cointegrate formation between pAt15955 and pTi15955 prior to Ti-driven conjugal transfer. To determine whether both plasmids were transferred during conjugation assays, pAt15955 transconjugants were screened for the presence of pTi15955 using octopine auxotrophy screening and Eckhardt gel electrophoresis (Figure 5 and Figure S7). As expected, donors possess pTi15955 and either pAt15955 (IB125) or pAt15955Δ270 (IB131), while the recipient (ERM52) is plasmidless (Figure 5). Transconjugants from either donor show a mix of plasmid profiles; some have both plasmids whereas others have only the pAt15955 or pAt15955Δ270 plasmids (Figure 5). The transconjugants without the Ti plasmid are unlikely to have arisen due to co-integrate formation, and for the pAt15955Δ270 plasmid the pTi15955-encoded conjugal machinery can compensate for the missing *trb* and *traI* genes. However, because the octopine-induced conjugation of pAt15955Δ270 appeared lower than pAt15955 (Figure 4), we tested whether the proportion of co-transfer of both plasmids was affected by the pAt15955 segment, and found that it does not impact the co-transfer frequency (Figure S7).

**Figure 5:**
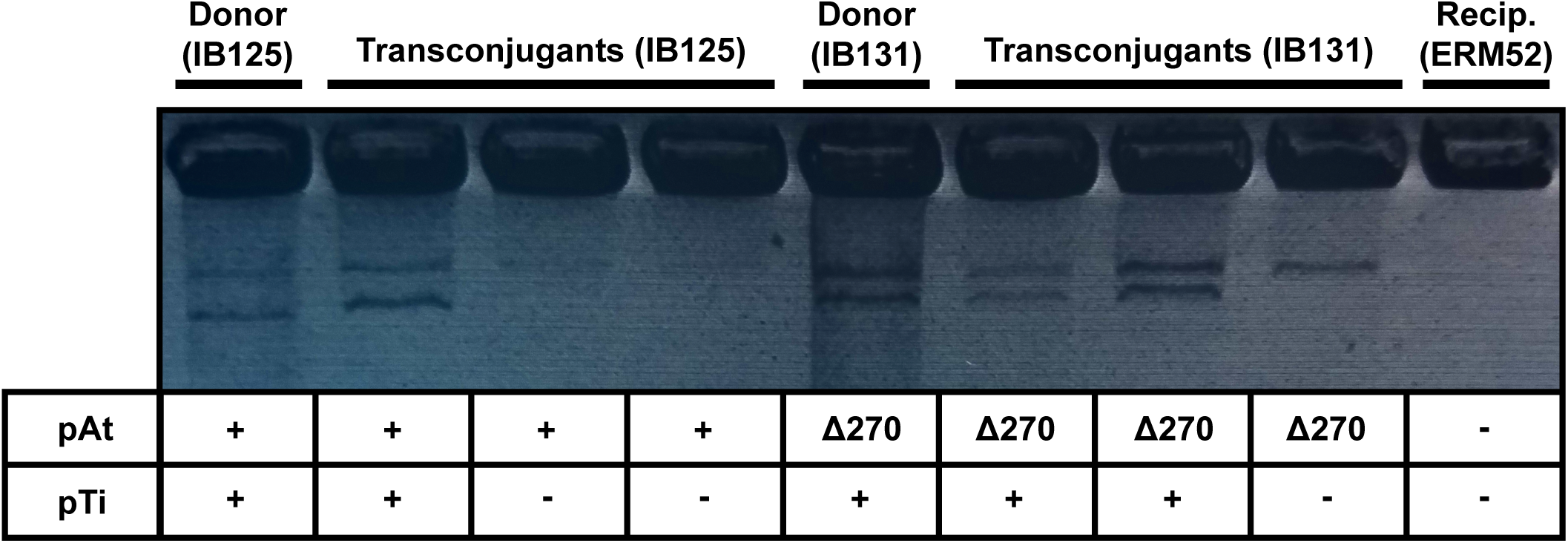
Co-transfer of pTi15955 in pAt15955 transconjugants in the presence of octopine. Plasmid profiles for subset of pAt15955 transconjugants from conjugation assay in Figure 4 were determined by Eckhardt gel electrophoresis. Shown are profiles for two donors (IB125 and IB131), recipient (ERM52), and three transconjugants from conjugation assays with either donor (IB125 or IB131). Indicated below gel image are presence (+) or absence (-) of pTi15955 and the presence of WT pAt15955 (+) or pAt15955Δ270 (Δ270).

### pAt15955Δ270 segment deletion has environment-dependent fitness consequences

To determine whether the pAt15955 segment imposes fitness consequences that may explain a propensity to lose the segment, competition experiments between isogenic strains that either carry or have lost the pAt15955 segment were performed under replete carbon, limiting carbon, and virulence-inducing conditions over nine serial passages (Figure 6). In replete carbon conditions, there was not a measurable fitness cost associated with the pAt15955Δ270 segment over the duration of the experiment. In limiting carbon conditions, there was a modest, measurable fitness effect of the segment that is similar to carriage costs determined for At plasmid deletions of a similar size in *A. tumefaciens* C58 (Morton *et al*. 2013). In virulence-inducing conditions, there was a significant fitness disadvantage to possessing the pAt15955 segment (Figure 6), suggesting that the presence of this region either increases general susceptibility to the stressful conditions of virulence-induction or potentiates the *vir* response under these conditions, both of which could result in a decrease of growth-based fitness as measured in this assay.

**Figure 6:**
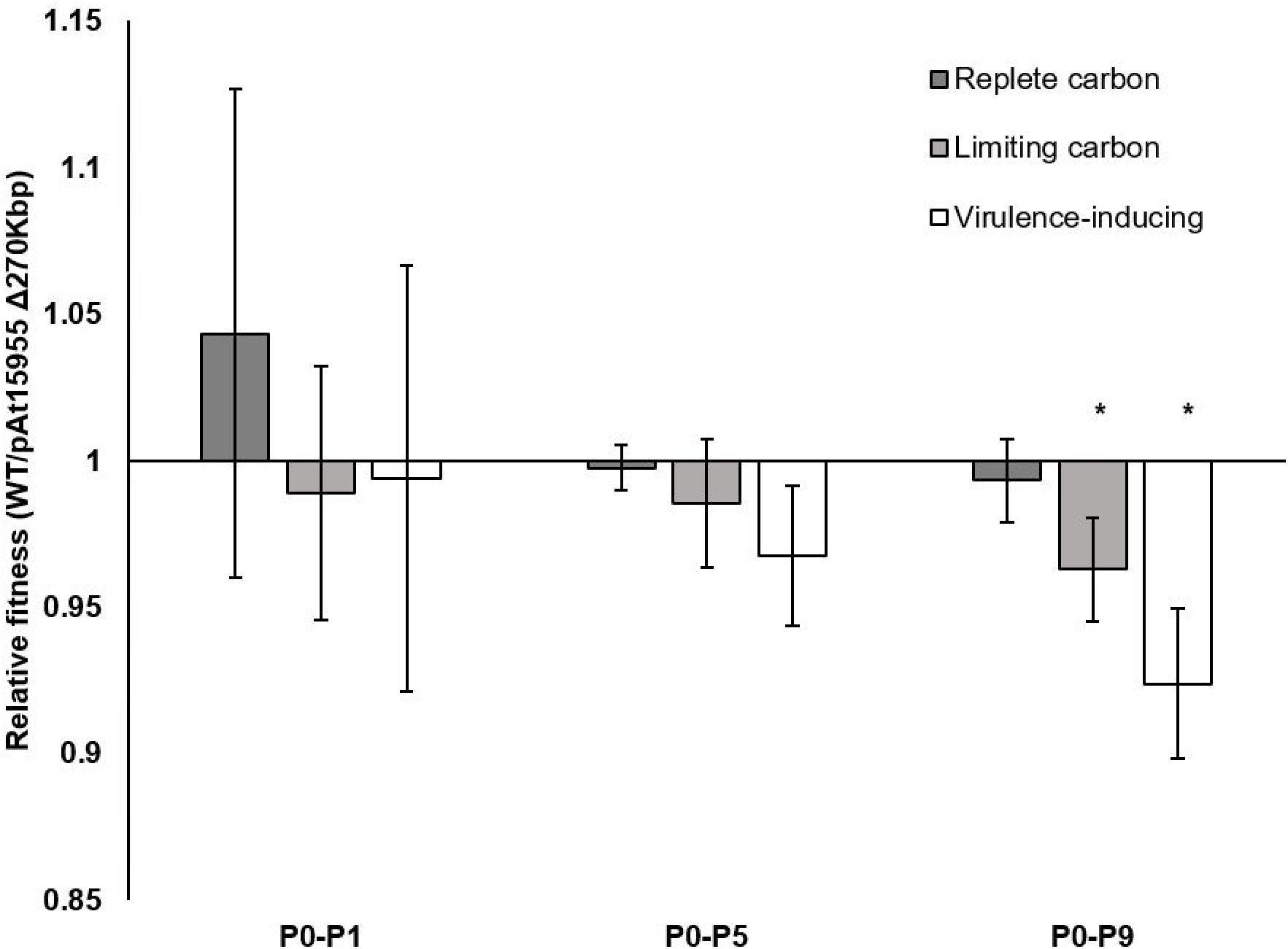
Absence of pAt15955 segment confers an environment-dependent fitness advantage. Relative fitness of WT *A. tumefaciens* 15955 and a strain isogenic for loss of the pAt15955 segment over one (P0-P1), five (P0-P5) and nine (P0-P9) serial passages in replete carbon (dark gray), limiting carbon (hatched), and virulence-inducing conditions (white). Bars represent 6 biological replicates, 3 each with inversely marked strains (Methods). Relative fitness calculated as the total number of doublings of each competitor over the indicated passing interval (Methods). Error bars shown are standard deviation. Asterisk indicates p-value < 0.05 in comparison to the expected value of 1.

## DISCUSSION

Members of the family *Rhizobiaceae* exhibit signs of replicon co-evolution (Slater *et al*. 2009; Lassalle *et al*. 2017; Wang *et al*. 2018) and cross-replicon interactions (Ronson *et al*. 1987; Heidelberg *et al*. 2000; Barnett *et al*. 2004; Bobik *et al*. 2006; Galardini *et al*. 2015; Pini *et al*. 2015; Ramachandran *et al*. 2017; diCenzo *et al*. 2018), despite remarkable strain-to-strain variation and the prevalence of large-scale rearrangements in secondary replicons (Orozco-Mosqueda Mdel *et al*. 2009; Mazur *et al*. 2011; Lopez-Guerrero *et al*. 2012; Morton *et al*. 2013; Galardini *et al*. 2015). *A. tumefaciens* is a member of this family and often carries a large proportion of its genomic DNA, including important niche-associated functions, on two large extrachromosomal megaplasmids, pAt and pTi (Goodner *et al*. 2001; Wood *et al*. 2001). Given the prevalence of both Ti and At plasmids within *A. tumefaciens* isolates and bioinformatic evidence suggesting co-evolution of these replicons (Lassalle *et al*. 2017), cross-replicon interactions are likely to exist and in part function to co-stabilize the replicons. To this effect, we have previously shown evidence of co-stability of both replicons in *A. tumefaciens* C58 (Morton *et al*. 2014), suggesting favorable interactions between both plasmids in some strains.

Here, we show that the process of curing the Ti plasmid in the octopine-type strain *A. tumefaciens* 15955 promotes a large deletion in the co-resident At plasmid, suggesting a discrete genetic link between both plasmids that may underlie co-replicon stability. Our observations suggested that the At plasmid might be limiting Ti curing. We found that Ti plasmid curing efficiency was slightly increased in derivatives missing the entire AtΔ270 fragment or ∼108 kb on the right half of this segment, suggesting that the genetic content in this region could impact overall Ti plasmid stability. However, these trends were not statistically robust and it seems unlikely that this potential stabilization effect explains the tight correlation between pTi curing and At plasmid segment loss that we report here.

The deletion event itself is mediated by a repetitive IS-element, IS66, that flanks the At segment and is required for its loss under curing conditions. IS66 was first identified on the Ti plasmid of a different octopine-type strain of *A. tumefaciens* (Machida *et al*. 1984) and is part of a larger family of IS-elements found within a variety of bacteria (Han *et al*. 2001; Gourbeyre *et al*. 2010). IS-elements are common vehicles for genome rearrangements and responsible for genome reduction in many bacterial taxa, including species of *Bordatella* (Parkhill *et al*. 2003; Siguier *et al*. 2006). Ti plasmid destabilization in the curing process could conceivably elicit loss of the At plasmid segment through transposition of both flanking IS66 elements in a mechanism similar to a composite transposon (Bellanger *et al*. 2014). However the observed retention of a single element at the site after deletion with distinct flanking duplications, and the tandem orientation of IS66 in full length pAt suggests homologous recombination as the mechanism for AtΔ270 segment loss, and hence should require RecA (Bell and Kowalczykowski 2016). Deletion of *recA* would clearly confirm whether homologous recombination is required for loss of the At plasmid segment. Whereas *recA* has been deleted previously in *A. tumefaciens* C58 (Farrand *et al*. 1989), our repeated attempts at generating a similar deletion in *A. tumefaciens* 15955 were unsuccessful (suggesting that the mutation is deleterious in 15955).

The IS66-mediated deletion reported here is different from another set of At plasmid large deletions we have described for *A. tumefaciens* C58, which appear to be mediated by small repeat sequences (Morton *et al*. 2013). The low degree of At plasmid synteny and the propensity of these plasmids to undergo large-scale rearrangements (Morton *et al*. 2013) suggests that the fitness benefit of gene content on the At plasmid may be highly environment-dependent. This is also true for the Ti plasmid, which imposes a heavy fitness burden upon exposure to virulence-inducing conditions (Platt *et al*. 2012a), as well as other organisms, including species of *Methylobacterium*, where experimental evolution resulted in an environment-dependent reduction of the accessory genome by ∼10% (Lee and Marx 2012).

Genomic stability can be maintained in variety of ways, including the reduction of costs associated with accessory genes (diCenzo and Finan 2017). For example, the stability of Ti plasmids is in part due to low carriage costs, low copy number, and extremely tight regulation of costly genes, including virulence functions and conjugal transfer (Pappas and Winans 2003; Pappas 2008; Platt *et al*. 2012a; 2012b; Morton *et al*. 2014). The current results have important consequences for our previous estimates of the low carriage costs of the pTi15955 because these experiments unknowingly involved strains that had incurred the pAt1599Δ270 kb deletion (Platt *et al*. 2012a). In our previous work, we concluded that under conditions which do not induce the Ti plasmid virulence genes and are not substantially expressed, there was a low but measurable carriage cost associated with pTi15955, while under virulence inducing conditions there was a large cost associated with pTi15955 (Platt *et al*. 2012a). The findings reported here show that the measurement of low, but significant basal carriage cost under non-inducing conditions reported in that paper is likely a combination of costs associated with the Ti plasmid and costs associated with the deleted At plasmid segment. The pAt15955Δ270 deletion alone results in a significant fitness gain under similar experimental conditions (Figure 6, p-values, <0.05, 6 independent replicates) and thus the carriage cost of the Ti plasmid reported in the previous paper is likely to be less than estimated (Platt *et al*. 2012a) and more similar to the low Ti plasmid carriage costs we determined for pTiC58 from the nopaline-type *A. tumefaciens* strain C58 (Morton et al. 2014).

The At plasmid deletion documented in this study had significant effects on overall fitness in an environment-dependent manner. While the segment conferred carriage costs similar to At plasmid deletions we have characterized previously (Morton *et al*. 2013), the burden of harboring the segment was significantly increased under virulence-inducing conditions. Virulence genes are induced by a combination of low phosphate and acidic pH, in addition to the presence of plant phenolics (Platt *et al*. 2012a), so these conditions are also highly stressful and may exacerbate the fitness cost of harboring this large segment of the megaplasmid. However, the increased cost of this segment of pAt under virulence induction also suggests a role for the At plasmid in the response of *A. tumefaciens* to the disease environment. This is consistent with our observations in *A. tumefaciens* C58, where the At plasmid was found to modulate virulence gene expression (Morton *et al*. 2013).

We identified additional interactions between these plasmids via their conjugal transfer functions. We observed that pAt15955 conjugation exhibits a basal level of conjugation that is further stimulated by the presence of octopine, the so-called conjugal opine for many octopine-type Ti plasmids (Klapwijk *et al*. 1978). The basal conjugation of pAt15955 in the absence of octopine is similar to that observed for the At plasmid of the nopaline-type *A. tumefaciens* strain C58, in which pAtC58 conjugates constitutively under laboratory conditions via its own conjugal transfer functions (Chen *et al*. 2002). The predicted pAtC58 conjugal pilus is encoded by the *avhB* genes (*<u>A</u>grobacterium* <u>v</u>irulence <u>h</u>omologue *vir<u>B</u>*), named as such due to their high homology to the pTiC58 plasmid-encoded VirB type IV secretion system which delivers T-DNA from *A. tumefaciens* to plants during infection. The *avhB* genes are more distantly similar to the pTiC58 conjugal pilus (*trb)* genes. In contrast, the pAt15955 *trb* genes within the AtΔ270 segment we describe here are more closely similar to the pTi15955 conjugal transfer (*tra/trb*) genes then they are to *virB* genes on pTi15955. The basal conjugation of pAt15955 is not affected by the presence of the pTi15955 plasmid, whereas the presence of pTiC58 decreases pAtC58 conjugation by 10-fold (Chen *et al*. 2002). However, activation of the pTiC58 *tra* genes was reported to stimulate pAtC58 conjugal transfer (Lang *et al*. 2013). This was hypothesized to proceed through activation of the *rctB* regulator of pAtC58 conjugal transfer functions. Interestingly, no *rctB*-type regulator was detected in the *A. tumefaciens* 15955 genome sequence, but pAt15955 instead encodes its own paralogues of the Ti plasmid quorum sensing control genes, *traR* and *traM* on the plasmid backbone, and the AHL synthese *traI* along with the *trb* genes, in the AtΔ270 segment. The TraR-TraI-TraM quorum sensing regulators on the Ti plasmid provide strict octopine and population density-dependent control of pTi conjugation for octopine-type Ti plasmids (Fuqua and Winans 1994; Fuqua *et al*. 1995; Fuqua and Winans 1996) and there is some evidence that in a close relative of 15955, the strain *A. tumefaciens* A6, these quorum sensing regulators may also regulate pAt conjugal transfer genes (Wang *et al*. 2014). The strong stimulation of pAt15955 by octopine, but only in the presence of the Ti plasmid, suggests that octopine may trigger these regulators to increase *tra* gene expression. Co-transfer of the At and Ti plasmid has been observed previously in the plant tumor environment (Lang *et al*. 2013), but we show here that it is directly related to octopine-induction of the Ti plasmid conjugation system and that both plasmid systems interact. Intriguingly, independent, octopine-inducible transfer of the pAt15955 plasmid that is missing the AtΔ270 segment which includes the *trb* and *traI* genes, only occurs in the presence of pTi15955, suggesting that pAt15955 may also be able to utilize the Ti *trb* functions for its own conjugation.

The genetic association between the At and Ti plasmids we have characterized suggests extensive cross-replicon interactions between these replicons that may underlie their overall stability in changing environments. As a large proportion of bacteria possess multiple large replicons, especially in *A. tumefaciens* and other members of *Rhizobiaceae*, similar interactions are likely to occur in those systems and may exert a profound influence on genomic stability. However, given the complex life histories associated with many of these bacteria, it will be challenging to identify the relevant environmental conditions which influence stability as well as the fitness costs associated with large-scale rearrangements.

## ACKNOWLEDGEMENTS

This work was supported by National Institutes of Health Grant [R01 GM092660 to C.F. and J.D.B] and the Indiana Faculty Research Support Program (FRSP). We thank James D. Bever (JDB) for serving as PI on the R01 grant. We also thank Grace Perkins for technical assistance in the plasmid curing studies, Dean Rowe-Magnus for reagents and advice for engineering large deletions using the Cre-*lox* system, and the Center for Genomics and Bioinformatics at Indiana University for their work preparing the Illumina libraries and operating the sequencing instrumentation.

SUPPLEMENTARY INFORMATION

**Figure S1:**
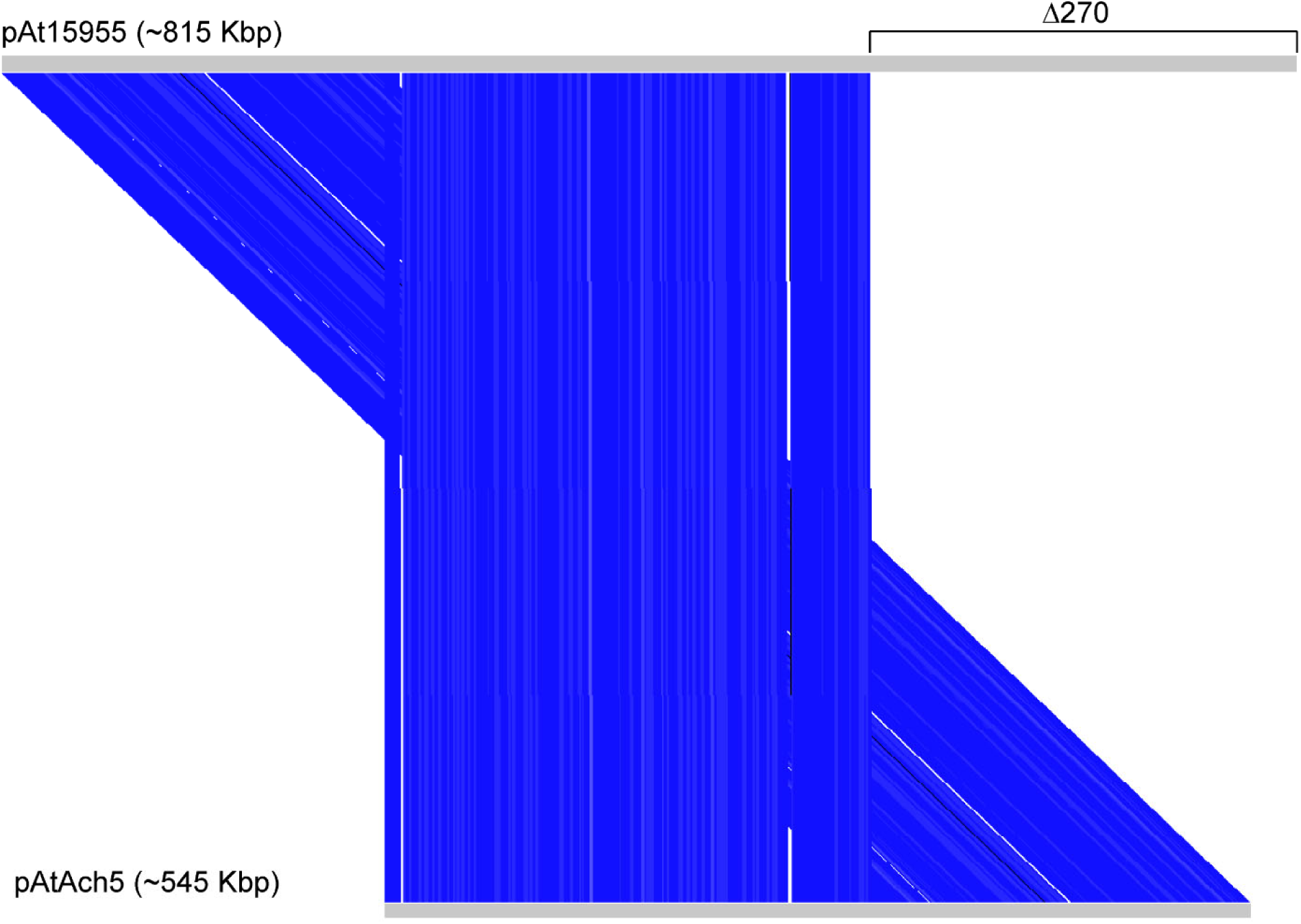
Analogous At plasmid deletion present in natural isolate, *A. tumefaciens* Ach5. Modified Artemis Comparison Tool (ACT) visualization of TBLASTX alignment between At plasmid sequences of *A. tumefaciens* 15955 and *A. tumefaciens* Ach5 indicates deletion in the *A. tumefaciens* Ach5 At plasmid (pAtAch5) at the precise location of the observed deletion in pAt15955. Gray bar depicts plasmid sequence. Blue connecting lines indicate stretches of homology >70% identity. Bracket indicates ∼270 Kbp deletion segment.

**Figure S2.**
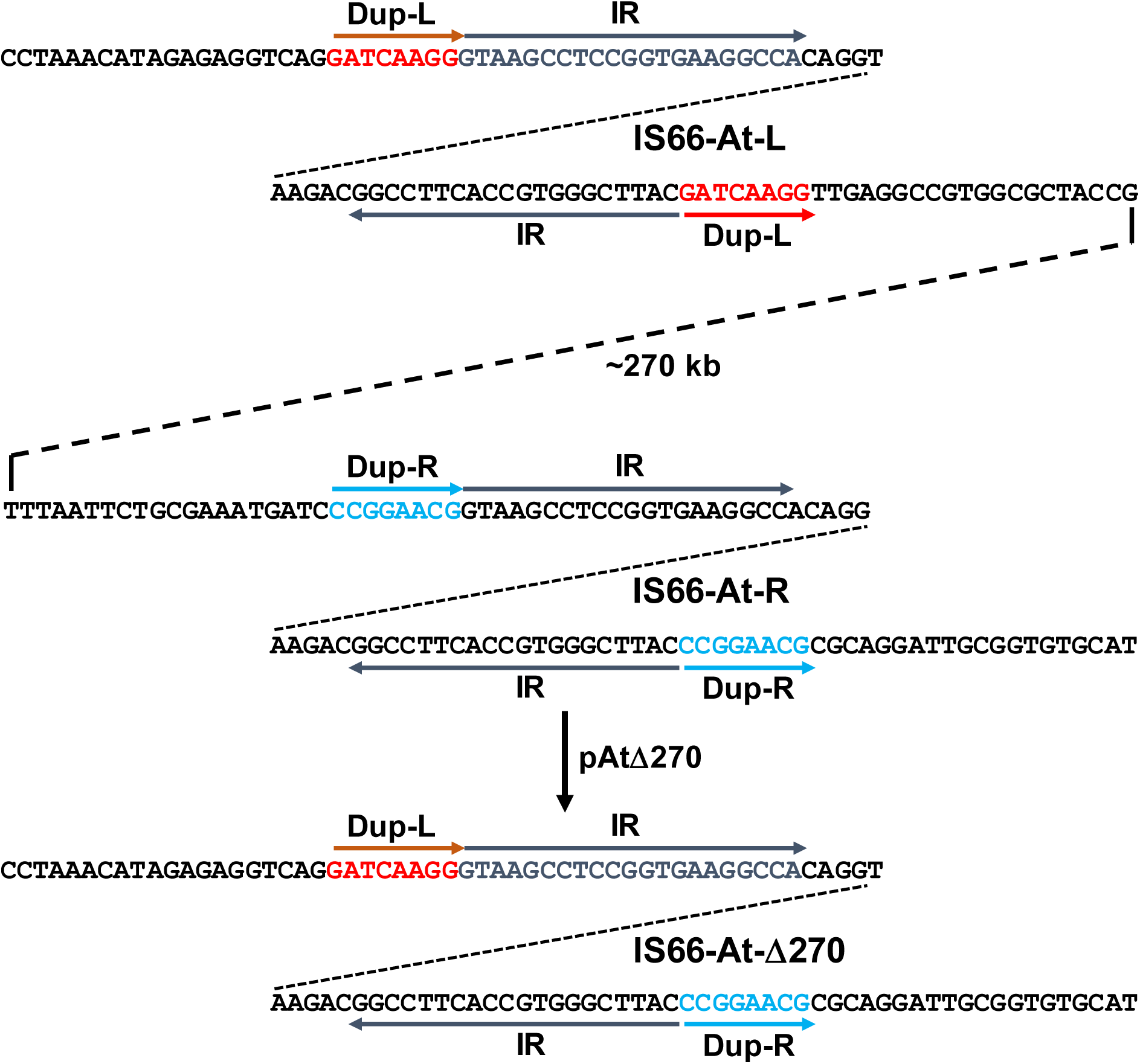
Flanking sequences from pAt15955 IS66 elements. IS66_At_L and IS66_At_R flank an ∼270kb segment of the At plasmid which is deleted upon pTi15955 plasmid curing. Light blue and red arrows labeled as Dup indicate the 8 bp duplication generated by IS66 insertion and arrows labeled IR indicate the inverted repeats that define the end of each IS66 element.

**Figure S3:**
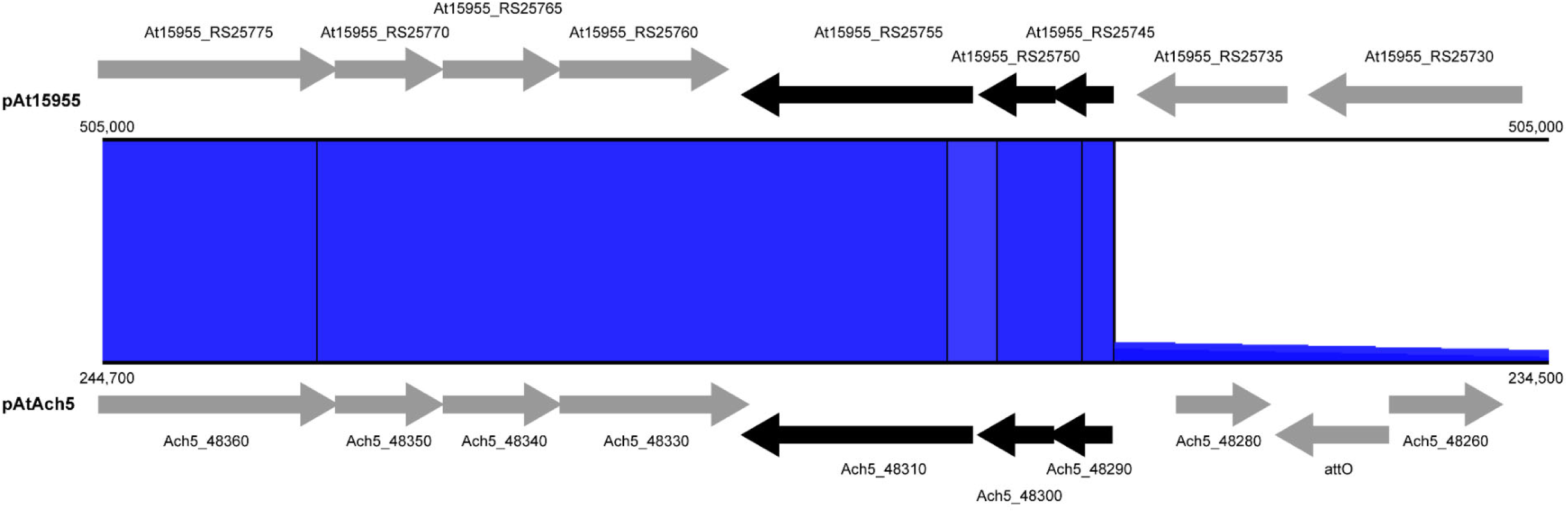
IS-element occurs at site of deletion in *A. tumefaciens* 15955 and *A. tumefaciens* Ach5. Modified ACT visualization of TBLASTX alignment between pAt15955 and pAtAch5 at deletion site identifies homologous IS-element at junction. Shown is alignment of junction site in pAtAch5 with one side of deletion interval in pAt15955 that still carries the Δ270 segment. Blue connecting lines indicate stretches of >90% identity

**Figure S4:**
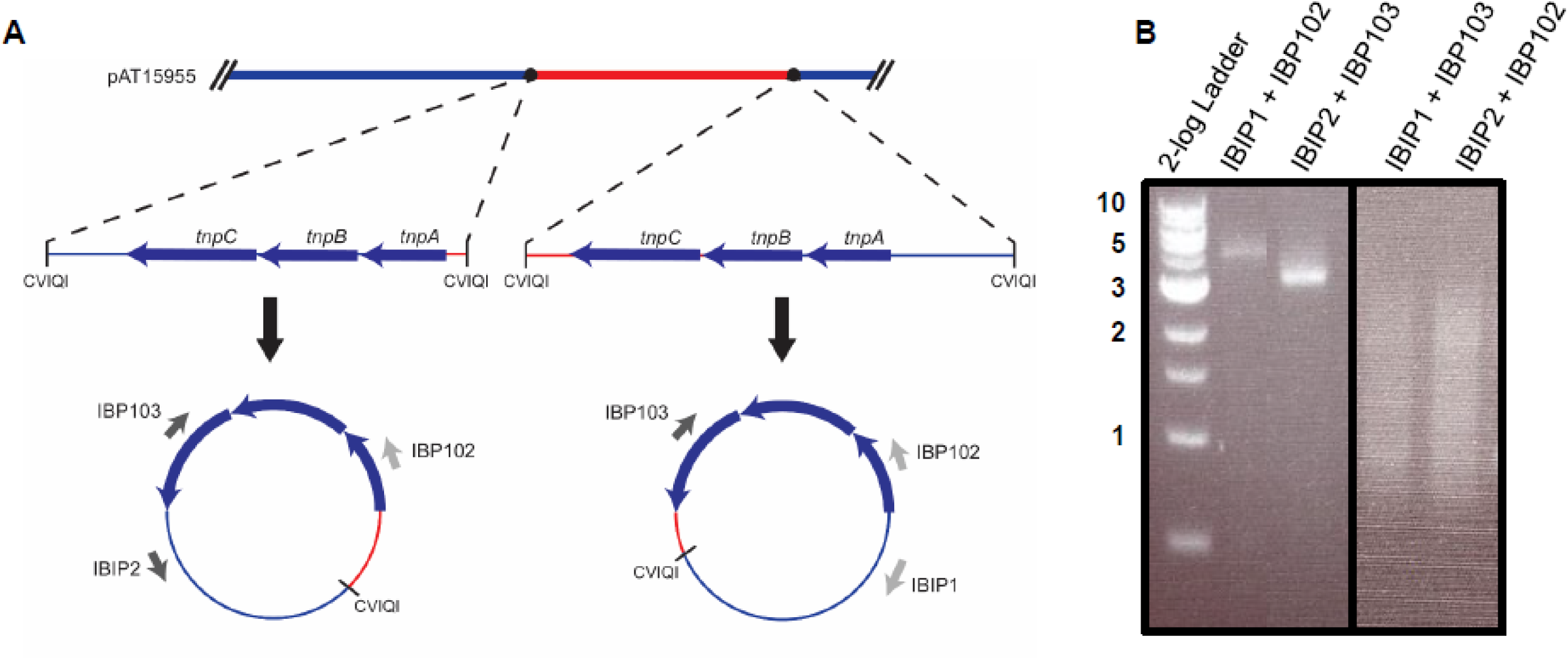
Inverse PCR approach confirms IS66 elements flank pAt15955 segment in a direct orientation. A) Schematic depicting inverse PCR design. gDNA from WT *A. tumefaciens* 15955 was digested with the restriction enzyme, CVQI, and subsequently ligated with T4 ligase. PCR analysis was then conducted on ligation products with sets of primers specific for either the IS66 element or flanking regions. Black circles or blue arrows represent IS66 elements. Red or blue lines indicate sequence within or without pAt15955 deletion segment, respectively. Larger, black arrows represent ligation step. Smaller, gray arrows represent PCR primers. Hash marks at end of replicon indicate continuation of sequence. Hashed lines indicate zoomed in depiction of region. B) Inverse PCR reactions performed on ligation products as depicted in A. Primer pairs used in each reaction are indicated. Ladder sizes shown in Kbp.

**Figure S5:**
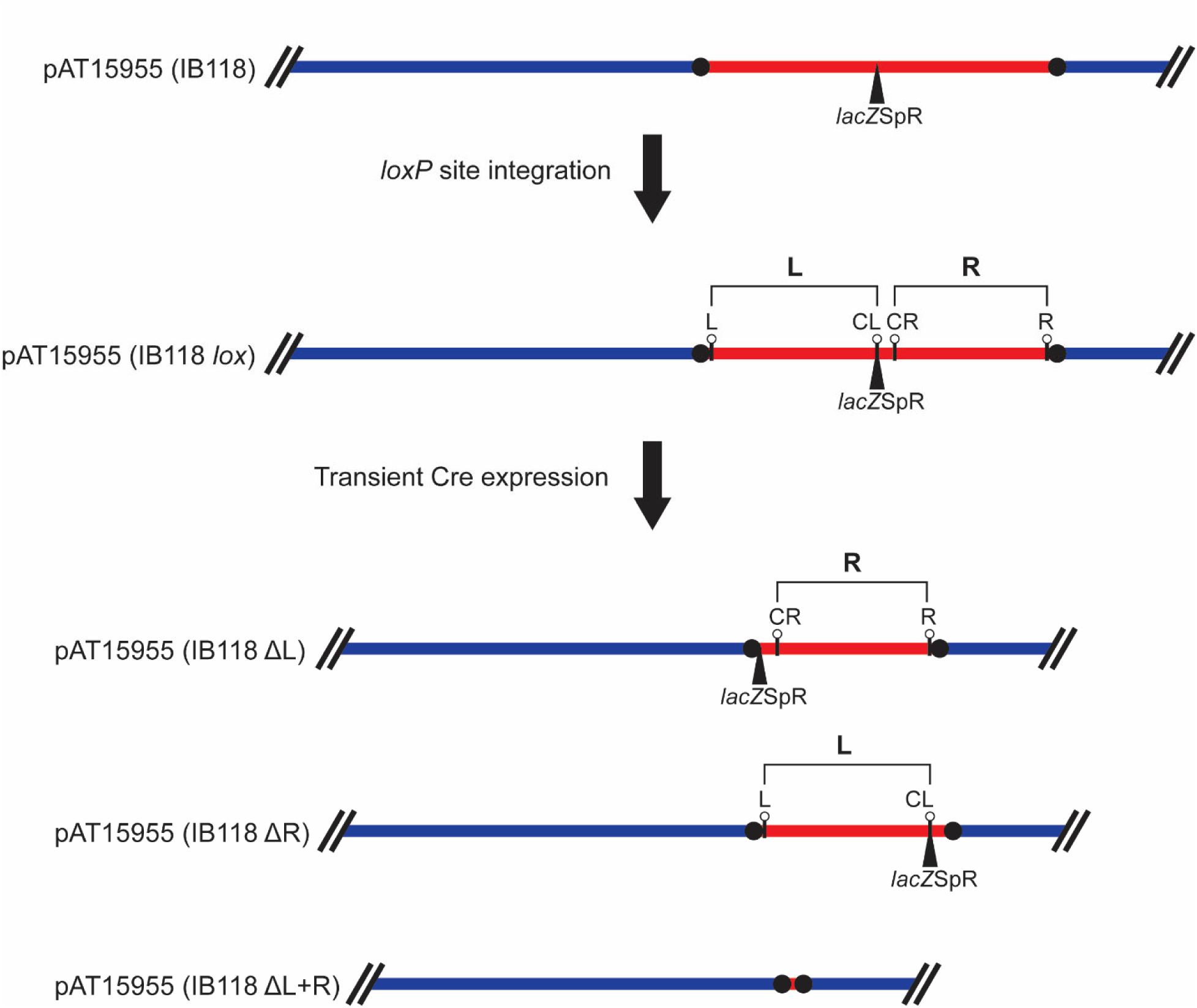
Schematic depicting Cre/*lox* approach for pAt15955 segment truncation. Cre/*lox* was utilized to truncated the pAt15955 segment within the reporter strain IB118. *loxP* sites were introduced sequentially into four locations within pAt15955 (denoted L, CL, CR, and R). All four sites are shown here (IB118 *lox*), but only two *loxP* sites were introduced in combination into three independent strains (see Methods, Table S1) to generate three variants upon transient Cre expression (IB118 ΔL, IB118 ΔR, and IB118 ΔL+R) that have been truncated for either the left (**L**) or right (**R**) half, or entire pAt15955 segment. Red or blue lines represent pAt15955 sequences within or without the pAt15955 deletion segment, respectively. Black circles indicate IS66 elements. *lacZ*Sp^R^ marker position is shown. Positions of *loxP* integration sites are shown

**Figure S6:**
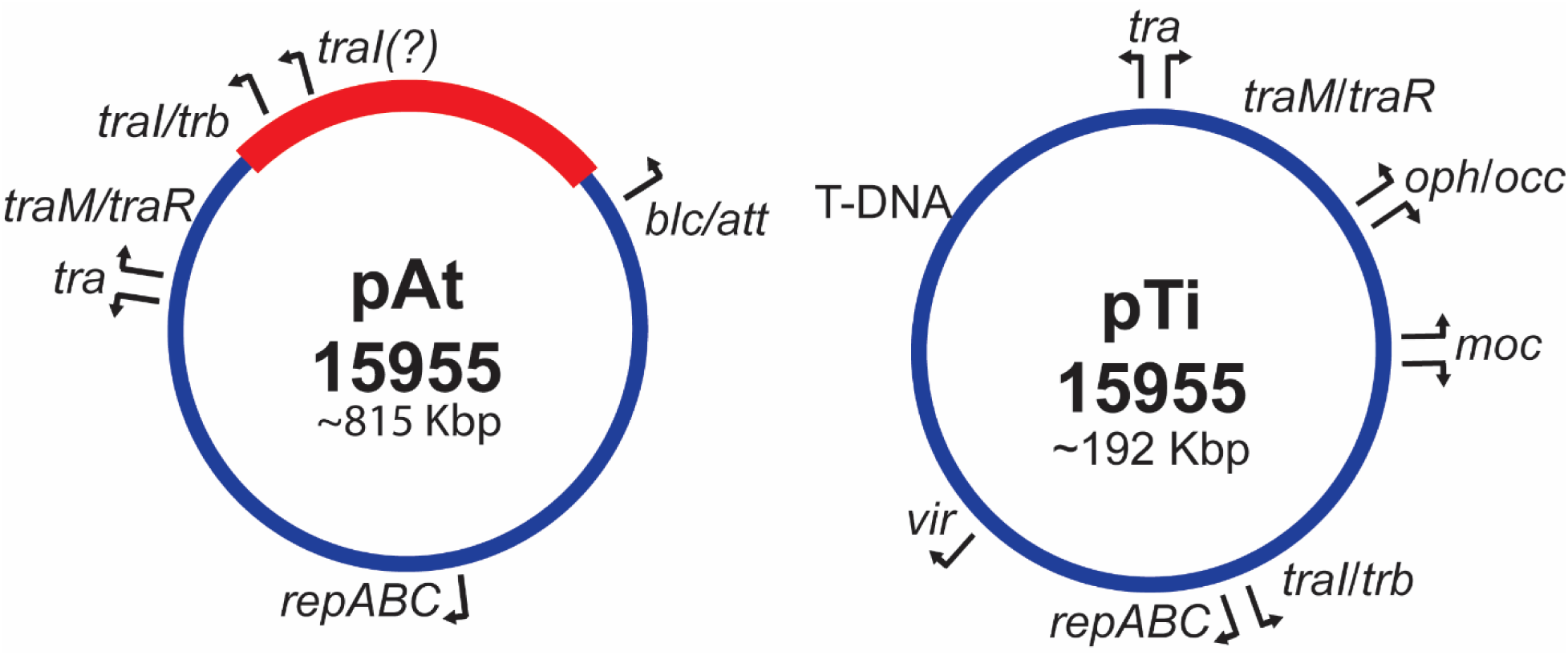
Organization of select genes within pAt15955 and pTi15955. Schematic showing placement of subset of operon locations within pAt15955 and pTi15955. The red segment in pAt15955 is the Δ270 region

**Figure S7:**
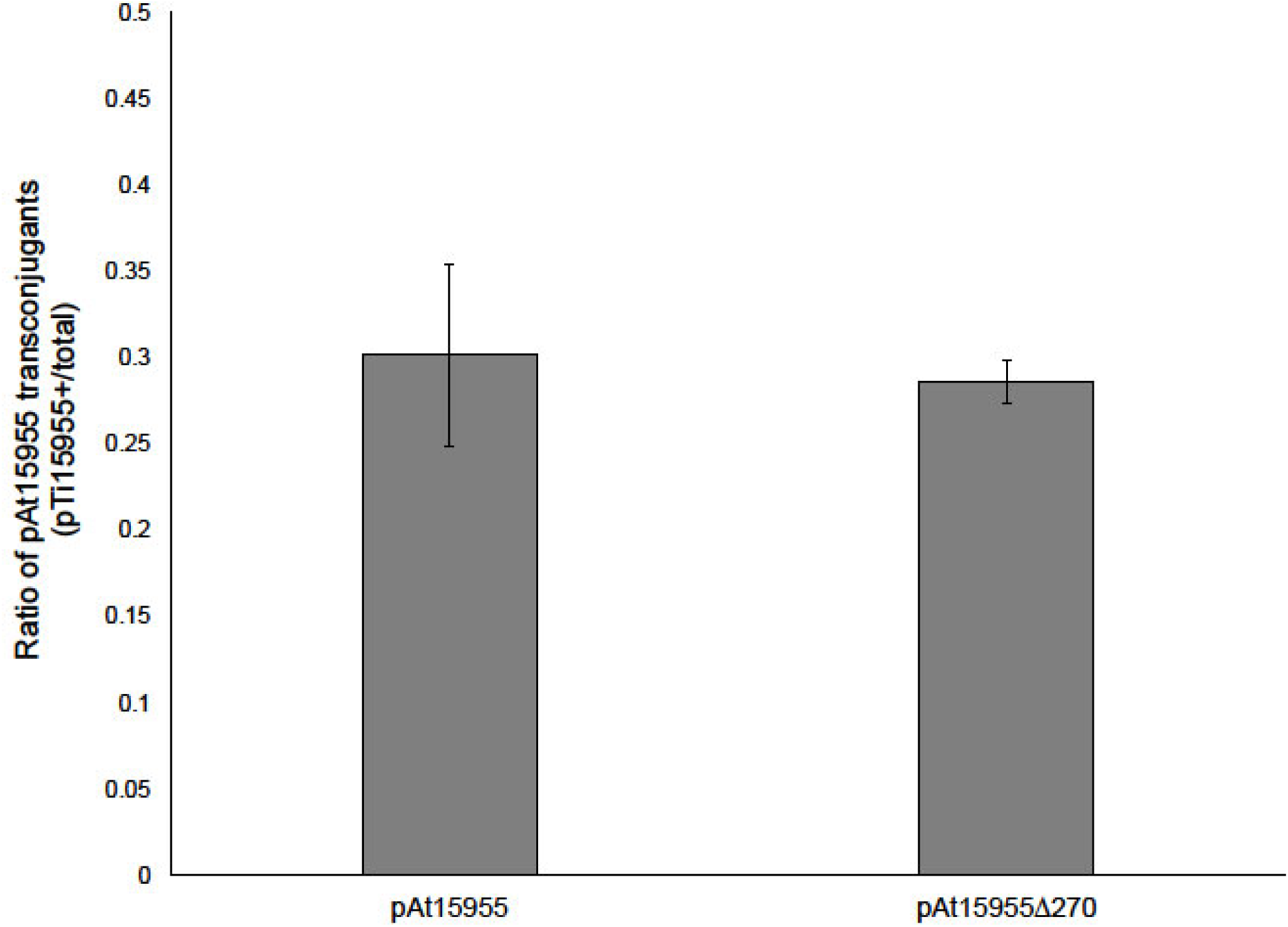
pAt15955 segment does not affect At and Ti plasmid co-transfer. Frequency of co-transfer between pAt15955 or pAt15955Δ270 and pTi15955 were determined by screening At plasmid transconjugants from IB125 or IB131 donor conjugations (Figure 4) for presence or absence of pTi15955 by octopine auxotrophy screening on ATO media (Methods). Co-transfer frequencies reported as the ratio of pTi15955 (+) to total transconjugants.

**Table S1:** Strains used in this study.

| Strain/plasmid | Genotype/Markers | Notes | Reference |
| --- | --- | --- | --- |
| ***E. coli*** |  |  |  |
| DH5α/λpir | λpir ; cloning strain |  | (Chiang et al. 2002) |
| S17-1/λpir | λpir ; Tra<sup>+</sup> , cloning host |  | (Kalogeraki et al. 1997) |
| ***A. tumefaciens* 15955** |  |  |  |
| WT | pAt15955;<br>pTi15955 |  |  |
| SA122 | pAt15955Δ270 | Cured of pTi15955<br>via chemical<br>methods | S.K. Farrand |
| IBE13A | pAt15955Δ270;<br>pTi15955 |  | This study |
| IB118 | pAt15955::lacZ/Sp<sup>R</sup>;<br>pTi15955::gusA/Gm<sup>R</sup> |  | This study |
| IB119 | Circ::Km<sup>R</sup> | Marked on circular<br>chromosome | This study |
| IB120 | Circ::Km<sup>R</sup>;<br>pAt15955Δ270;<br>pTi15955 | IBE13A marked on<br>circular<br>chromosome | This study |
| IB123 | pAt15955;<br>pTi15955::gusA/Gm<sup>R</sup> |  | This study |
| IB118ΔIS66 | pAt15955::lacZ/Sp<sup>R</sup>;<br>pTi15955::gusA/Gm<sup>R</sup>;<br>ΔIS66 | IB118 that has been<br>deleted flanking<br>IS66 on pAt15955 | This study |
| IB124 | pAt15955Δ270 | Cured of pTi15955<br>via incompatibility | This study |
| IBJUN138 | pAt15955 | pTi15955-cured<br>derivative with intact<br>pAt15955 | This study |
| IB125 | pAt15955::Km<sup>R</sup>;<br>pTi15955 |  | This study |
| IB126 | Circ::Sp<sup>R</sup>;<br>pAt15955;<br>pTi15955::gusA/Gm<sup>R</sup> | IB123 marked on<br>circular<br>chromosome | This study |
| IB127 | Circ::Sp<sup>R</sup>;<br>pAt15955Δ270 | IB124 marked on<br>circular<br>chromosome | This study |
| IB128 | Circ::Sp<sup>R</sup>;<br>pAt15955 | IBJUN138 marked<br>on circular<br>chromosome | This study |
| IB129 | Circ::Sp<sup>R</sup>;<br>pAt15955Δ270;<br>pTi15955::gusA/Gm<sup>R</sup> | IB127 with pTi15955<br>reintroduced through<br>mating with IB123 | This study |
| IB130 | Circ::Sp<sup>R</sup>;<br>pAt15955;<br>pTi15955::gusA/Gm<sup>R</sup> | IB128 with pTi15955<br>reintroduced through<br>mating with IB123 | This study |
| IB118 *loxL* | pAt15955/*loxL*::*lacZ*/Sp<sup>R</sup>;<br>pTi15955::gus/Gm<sup>R</sup> | IB118 with two *loxP*<br>sites introduced<br>within left half (**L**) of<br>pAt15955 deletion<br>interval | This study |
| IB118 *loxR* | pAt15955/*loxR*::*lacZ*/Sp<sup>R</sup>;<br>pTi15955::gusA/Gm<sup>R</sup> | IB118 with two *loxP*<br>sites introduced<br>within right half (**R**)<br>of pAt15955 deletion<br>interval | This study |
| IB118 *loxL+R* | pAt15955/*loxL+R*::*lacZ*/Sp<sup>R</sup>;<br>pTi15955::gusA/Gm<sup>R</sup> | IB118 with two *loxP*<br>sites introduced<br>within pAt15955<br>deletion interval | This study |
| IB118 ΔL | pAt15955ΔL::*lacZ*/Sp<sup>R</sup>;<br>pTi15955::gusA/Gm<sup>R</sup> | IB118 *loxL* with<br>truncation of left half<br>of pAt15955<br>segment (~130Kbp)<br>via Cre | This study |
| IB118 ΔR | pAt15955ΔR::*lacZ*/Sp<sup>R</sup>;<br>pTi15955::gusA/Gm<sup>R</sup> | IB118 *loxR* with<br>truncation of right<br>half of pAt15955<br>segment (~130Kbp)<br>via Cre | This study |
| IB118 ΔL+R | pAt15955ΔL+R;<br>pTi15955::gusA/Gm<sup>R</sup> | IB118 *loxL+R* with<br>truncation of entire<br>pAt15955 segment<br>(~270Kbp) via Cre—<br>removing 1234 *lacZ*<br>Sp<sup>R</sup> marker | This study |
| IB131 | pAt15955Δ270::Km<sup>R</sup><br>pTi15955 | IBE13A with Km<sup>R</sup><br>marker on<br>pAt15955, outside of<br>deletion interval | This study |
| IB132 | pAt15955Δ270::Km<sup>R</sup> | IB124 with Km<sup>R</sup> marker on pAt15955, outside of deletion interval | This study |
| IB133 | pAt15955::Km<sup>R</sup> | IBJUN138 with Km<sup>R</sup> marker on pAt15955, outside of deletion interval | This study |
| ERM52 | Circ::Sp<sup>R</sup> | *A. tumefaciens* C58 derivative, Plasmidless Ti- Ai- | (Morton et al. 2014) |

**Table S2:**
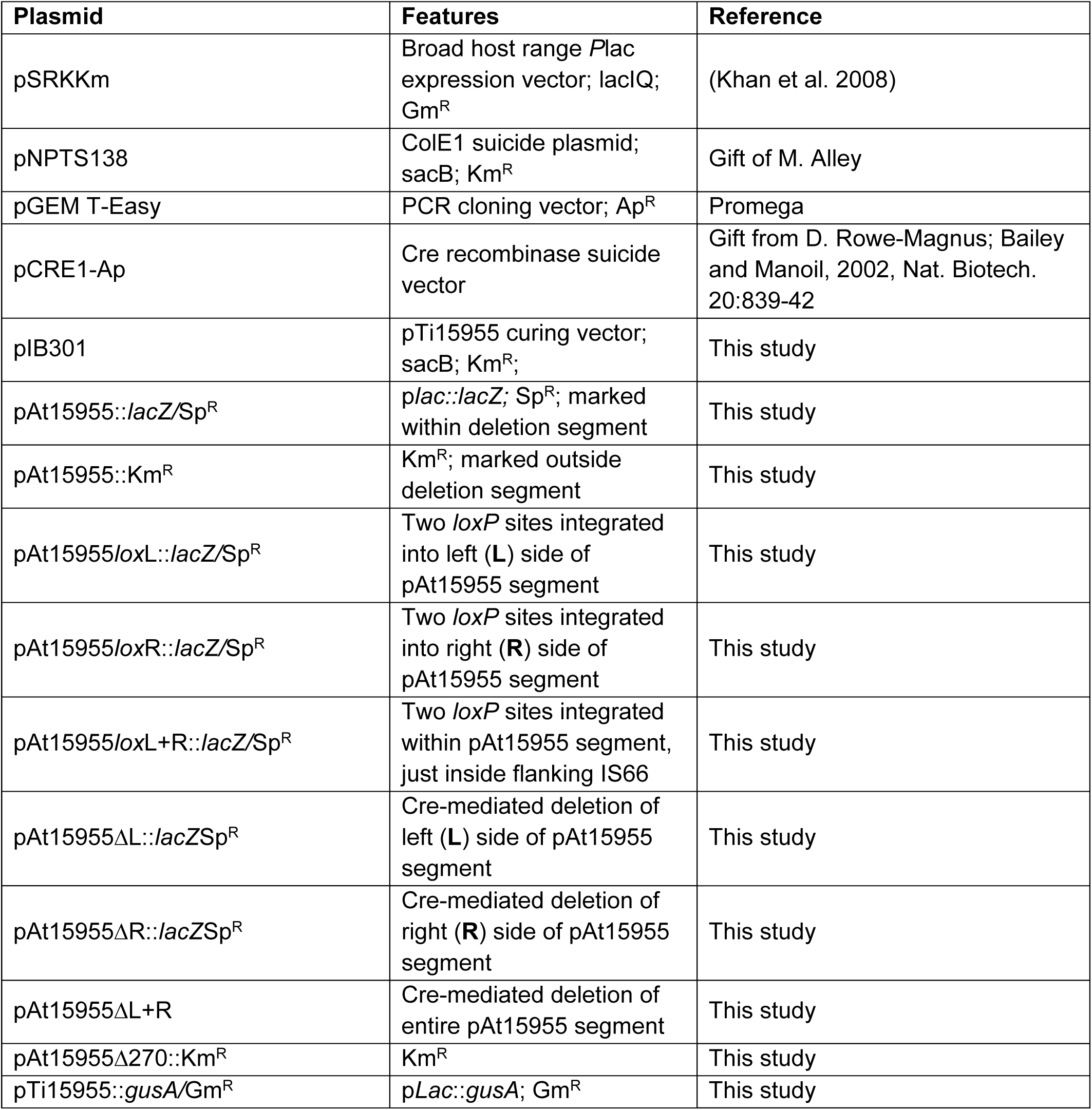
Plasmids used in this study.

**Table S3:**
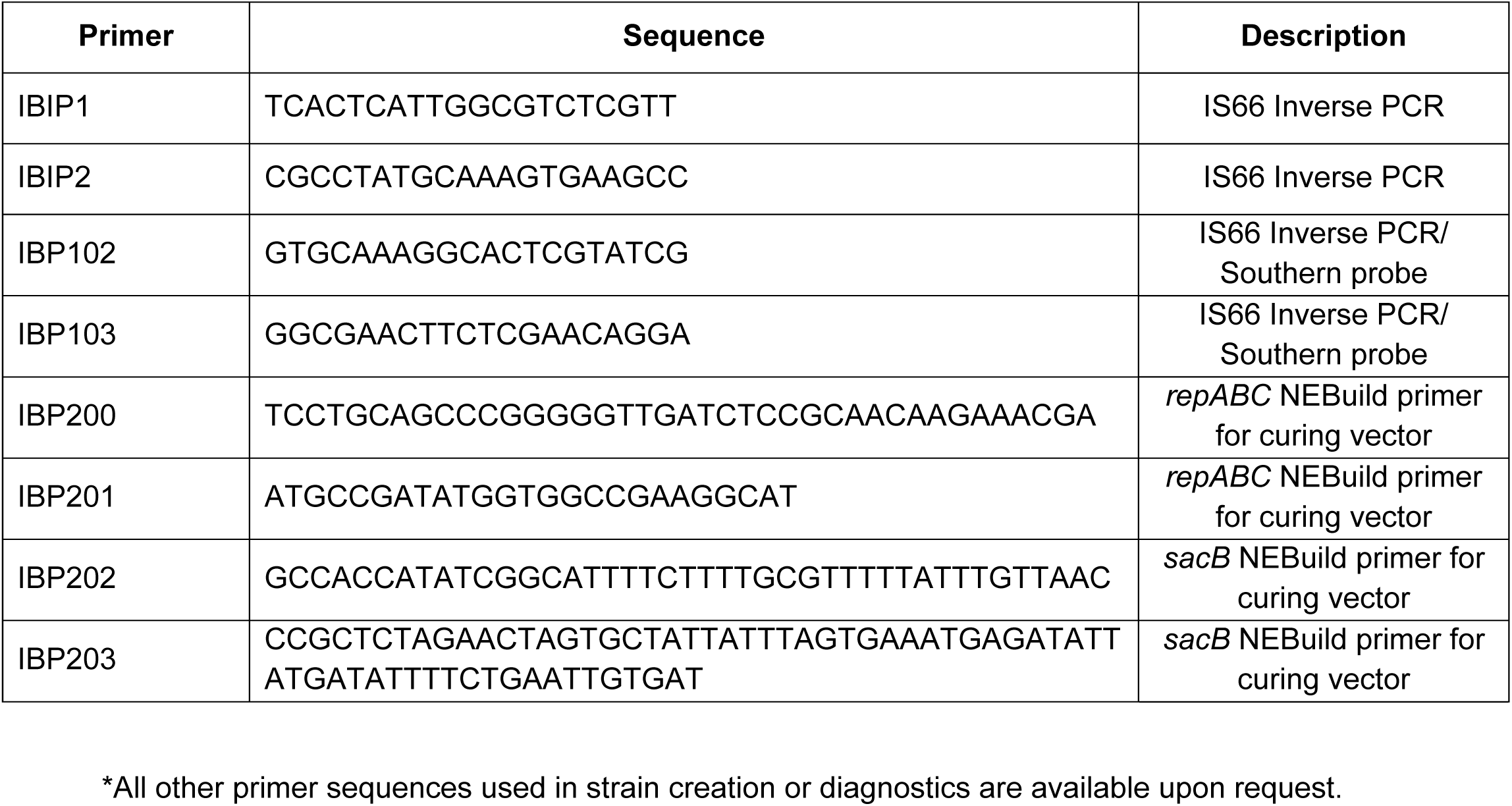
Primers used in this study*.

